# Increasing the accuracy of single-molecule data analysis using tMAVEN

**DOI:** 10.1101/2023.08.15.553409

**Authors:** Anjali R. Verma, Korak Kumar Ray, Maya Bodick, Colin D. Kinz-Thompson, Ruben L. Gonzalez

**Affiliations:** Department of Chemistry, Columbia University, New York, NY 10027 USA; Department of Chemistry, Rutgers University-Newark, Newark, NJ 07102 USA

## Abstract

Time-dependent single-molecule experiments contain rich kinetic information about the functional dynamics of biomolecules. A key step in extracting this information is the application of kinetic models, such as hidden Markov models (HMMs), which characterize the molecular mechanism governing the experimental system. Unfortunately, researchers rarely know the physico-chemical details of this molecular mechanism *a priori*, which raises questions about how to select the most appropriate kinetic model for a given single-molecule dataset and what consequences arise if the wrong model is chosen. To address these questions, we have developed and used time-series <u>M</u>odeling, <u>A</u>nalysis, and <u>V</u>isualization <u>EN</u>vironment (tMAVEN), a comprehensive, open-source, and extensible software platform. tMAVEN can perform each step of the single-molecule analysis pipeline, from pre-processing to kinetic modeling to plotting, and has been designed to enable the analysis of a single-molecule dataset with multiple types of kinetic models. Using tMAVEN, we have systematically investigated mismatches between kinetic models and molecular mechanisms by analyzing simulated examples of prototypical single-molecule datasets exhibiting common experimental complications, such as molecular heterogeneity, with a series of different types of HMMs. Our results show that no single kinetic modeling strategy is mathematically appropriate for all experimental contexts. Indeed, HMMs only correctly capture the underlying molecular mechanism in the simplest of cases. As such, researchers must modify HMMs using physico-chemical principles to avoid the risk of missing the significant biological and biophysical insights into molecular heterogeneity that their experiments provide. By enabling the facile, side-by-side application of multiple types of kinetic models to individual single-molecule datasets, tMAVEN allows researchers to carefully tailor their modeling approach to match the complexity of the underlying biomolecular dynamics and increase the accuracy of their single-molecule data analyses.

**Statement of Significance:** The power of time-dependent single-molecule biophysical experiments lies in their ability to uncover the molecular mechanisms governing experimental systems by computationally applying kinetic models to the data. While many software solutions have been developed to estimate the optimal parameters of such models, the results reported here show that the models themselves are often inherently mismatched with the molecular mechanisms they are being used to analyze. To investigate these mismatches and demonstrate how to best model the kinetics of a molecular mechanism, we have used time-series <u>M</u>odeling, <u>A</u>nalysis, and <u>V</u>isualization <u>EN</u>vironment (tMAVEN), an open-source software platform we have developed that, among other features, enables the analysis of single-molecule datasets using different kinetic models within a single, extensible, and customizable pipeline.

## Introduction

Single-molecule kinetics experiments have revolutionized our understanding of the dynamics of biomolecules and, consequently, the mechanisms of biomolecular function (1, 2). These techniques provide a uniquely detailed view of molecular mechanisms relative to more traditional ‘bulk’ techniques where mechanistic details, such as transient intermediates or rare events, are easily masked by macroscopic ensemble averaging. Furthermore, by observing the motions of an individual molecule through time, single-molecule kinetics experiments bypass the difficulties of biochemically synchronizing a stochastic biomolecular process with asynchronous dynamics (3). Altogether, the advantages conferred by single-molecule techniques have expanded both the set of biological systems whose kinetics can be investigated and the complexity of biomolecular dynamics that may be experimentally probed.

Despite the advantages of single-molecule techniques, the challenges of extracting mechanistic information from the associated experimental data (4) have limited their widespread application. Fortunately, many analysis approaches have been developed over the last few decades to address this problem (4, 5). In particular, hidden Markov models (HMMs) (6) have emerged as a popular method to describe the latent dynamics of a biological system by analyzing the time-dependent readout of a chosen experimental single-molecule signal (*e.g*., fluorescence intensity, spatial position, end-to-end distance, electric current, *etc*.) (7–21). In this context, HMMs are used to mathematically describe the transitions between relatively stable signal values (*i.e*., signal states) that are often observed in single-molecule experiments, thereby directly extracting the underlying kinetics of the molecular system from the experimental data.

A major consideration when analyzing single-molecule data with models like HMMs is the completeness of the kinetic information present in a signal *vs*. time trajectory. Each trajectory reports on the dynamic behavior of an individual molecule, thus the amount of molecular information that can be extracted from any one trajectory is limited (4)—particularly for techniques that rely on fluorescence, wherein the photophysical processes of the fluorophores severely restrict the length of the trajectory (22). In such cases, researchers often use their knowledge of the underlying physico-chemical properties of the biomolecular system to invoke the ergodic hypothesis (23) and model data from multiple trajectories in aggregate (*e.g*., as in Refs. (24–27)). Thus, instead of modeling the kinetic behavior of an individual molecule, these analyses infer the behavior of a mesoscopic, homogenous ensemble consisting of hundreds to thousands of molecules that are assumed to be identical.

In reality, however, experimental ensembles of molecules are never identical and always exhibit some amount of heterogeneity (28–31). These heterogeneities may be intrinsic to the nature of the biomolecule, such as the presence of several subpopulations of molecules within the experimental ensemble, or the presence of multiple molecular processes occurring over a range of timescales (Fig. 1). In other cases, heterogeneities may be artifacts of the specific experimental technique employed (*e.g*., interactions of the biomolecule with a surface in surface-tethered single-molecule experimental modalities), which must still be accounted for to avoid obscuring the underlying mechanistic details. Regardless of the source, such heterogeneities reduce the apparent complexity of the biomolecular system under investigation by collapsing distinct kinetic processes into the same signal state. This creates a mismatch between the ‘true’ underlying molecular mechanism reported by the experiment (Fig. 1) and the kinetic model that a researcher might choose for analysis based on the observed signal (Fig. 2) (*i.e*., a model-mechanism mismatch). Furthermore, while prior knowledge of the type or amount of heterogeneity present in an experimental dataset may guide the choice of kinetic model, such knowledge is not always readily available. Faced with a novel single-molecule dataset and a range of different kinetic models, the choice of which kinetic model to use is non-trivial and researchers are likely to incorrectly select a mismatched model. While we naturally expect the application of a mismatched model to affect the accuracy of the subsequent single-molecule analysis, the exact effects of such mismatches on the inferred mechanistic details are unknown.

**Figure 1.**
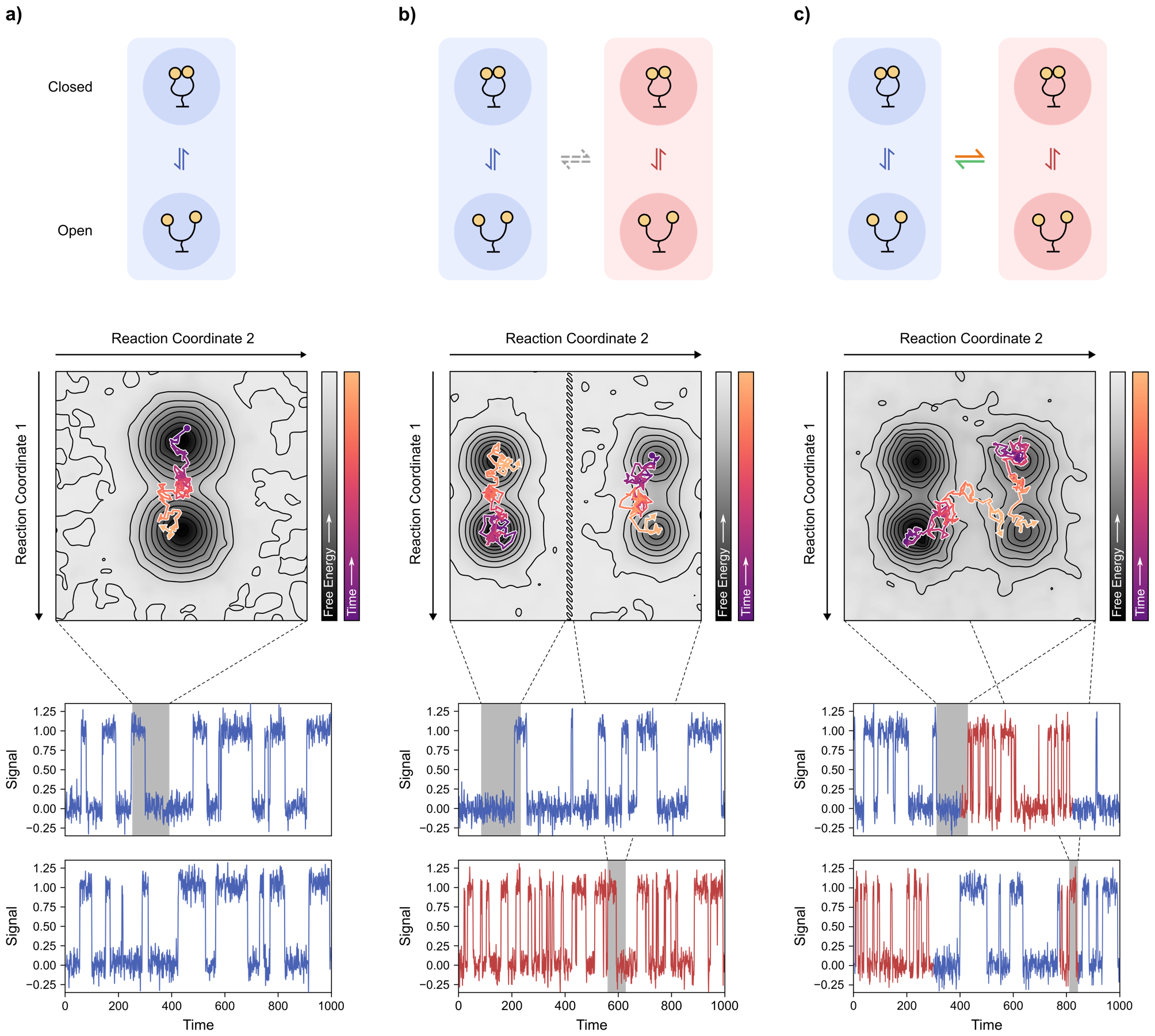
Molecular mechanisms and their corresponding single-molecule signal vs. time trajectories. (top) Schematic of the molecular mechanism, (middle) the corresponding conformational free-energy landscape, and (bottom) single-molecule trajectories that capture changes in signal for Reaction Coordinate 1 for **(a)** homogeneous, **(b)** statically heterogeneous, and **(c)** dynamically heterogeneous biomolecular systems. Simulated random walkers on the conformational free-energy landscape, starting at circles and ending at arrows, show hypothetical individual molecules undergoing transitions that correspond to the grey areas of the single-molecule trajectories. For the heterogeneous cases, blue and red correspond respectively to slow and fast transitioning subpopulations (for static) and phases (for dynamic), which are differentiated along Reaction Coordinate 2. A discontinuity (hatched line) is shown in the landscape for **(b)** to signify the lack of allowed transition along Reaction Coordinate 2 in this case.

**Figure 2.**
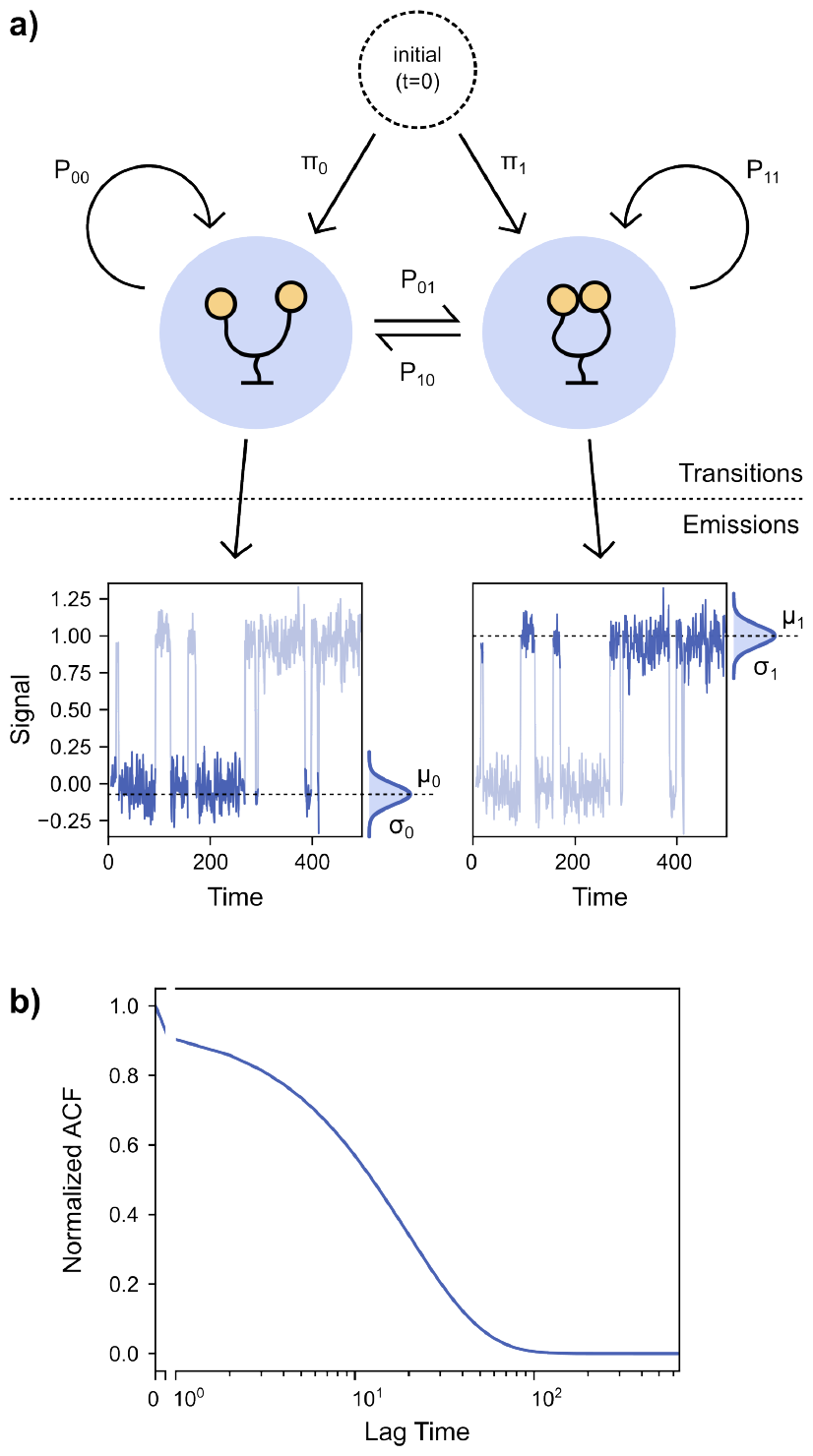
Schematic diagram of a kinetic model. **(a)** A schematic diagram of a two-state HMM showing the separation between the transition DoFs comprised of the initial probabilities and the transition probabilities, and the emission DoFs comprised of the emission probability distributions. **(b)** The normalized ACF corresponding to the HMM in **(a)** expresses all the dynamics of the kinetic model from both the transitions and the emissions in a single analytical form.

In this work, we investigate the effects of model-mechanism mismatches using a range of simulated datasets representative of commonly encountered types of heterogeneity in single-molecule experiments (Fig. S1 and Supplementary Information). To perform this analysis, we used a comprehensive, open-source, and extensible analysis platform we have developed, called time-series <u>M</u>odeling, <u>A</u>nalysis, and <u>V</u>isualization <u>EN</u>vironment (tMAVEN). tMAVEN is a software platform that, in addition to enabling the pre-processing of single-molecule time-series data, facilitates the interchangeable application of multiple, distinct kinetic models to the same single-molecule dataset. Beyond the above features, tMAVEN has also been designed to generate reproducible, publication-quality visualizations of experimental data and various modeling outcomes—all within a single computational pipeline.

Utilizing the broad range of modeling approaches implemented in tMAVEN, we show that only by exactly matching the number of free parameters (*i.e*., the complexity) of the underlying molecular mechanism does a kinetic model accurately infer the often-heterogeneous biomolecular dynamics under investigation. As such, no kinetic modeling strategy is universally appropriate across all experimental contexts. Moreover, this requirement of mechanism matching is separate from, and more fundamental than, questions about the performance of specific algorithmic or software implementations of a model (32), or the strategies used to optimize the parameters of such a model (*e.g*., maximum likelihood estimation, Bayesian inference, neural networks, *etc*.) (4). In the absence of a universal kinetic model, our study of where and how kinetic models fail in capturing the dynamics of heterogeneous, mesoscopic ensembles of single molecules can aid researchers in determining the optimal approach to quantifying the molecular details of their biomolecular system. Taken together with the capabilities that tMAVEN provides for applying multiple types of kinetic models to individual single-molecule datasets, our investigation facilitates the highly context-dependent kinetic modeling that is required for accurate single-molecule data analysis and, consequently, maximum extraction of biochemical and biophysical insight from single-molecule experiments.

## Theory

### Molecular mechanisms can be mapped onto a conformational free-energy landscape

The molecular mechanism through which a biomolecule undergoes a structural rearrangement and/or binding and dissociation process may be explained using the theoretical framework of a conformational free-energy landscape (33). A conformational free-energy landscape is a low-dimensional projection of the conformational space of a biomolecule onto the relevant reaction coordinates that characterize the specific process of interest, where each point represents the free energy of a specific set of biomolecular conformations. Thus, mapping a biomolecular system onto its corresponding conformational free-energy landscape provides a representation of the molecular mechanism underlying the dynamics of the system. Note that since most single-molecule techniques probe conformational changes, we have restricted our usage to conformational free-energy landscapes, but our discussion generalizes to other types of free-energy landscapes (*e.g*., chemical free-energy landscapes that represent chemical transformations).

In this work, we have considered a hypothetical biomolecule that exists in two conformational states (‘open’ and ‘closed’) with transitions between these states that are governed by first-order kinetics (Fig. 1a). On the corresponding conformational free-energy landscape, the open and closed conformations are represented by two minima or ‘wells.’ The open and closed wells are separated by a region of higher free energy that serves as a barrier to the transition between these two conformational states along the relevant reaction coordinate (*i.e*., the transition state). The height of this barrier (*i.e*., the difference in free energy between the minima of the well and the maxima of the transition state) is a single independent parameter that controls the rate of transitions between the open and closed states (34). For our hypothetical biomolecule, the heights of the barriers for the open→closed and closed→open reactions comprise two independent parameters, representing two degrees of freedom (DoFs), that together define the molecular mechanism involved in the open⇌closed equilibrium. It is worth noting that these two DoFs may be parameterized in different, but equivalent, manners (*e.g*., two barrier heights, the height of one barrier and the free-energy difference between the states, two rate constants, the equilibrium constant and the relaxation lifetime, *etc*.), but that two independent parameters are always required to describe this particular equilibrium. Together, these two independent parameters comprise the ‘mechanistic’ DoFs for our two-state system.

A final consideration in the mapping of a molecular mechanism onto a conformational free-energy landscape is how these conformational states are observed in a particular single-molecule experiment. In the ideal case above, we have represented these conformational states as smooth minima on the conformational free-energy landscape that each generate a distinct signal. Real biomolecules, however, exist in a hierarchy of conformational states (*i.e*., wells within wells) and consequently traverse ‘rugged’ conformational free energy landscapes (33, 35, 36). Depending upon the experimental time resolution used to probe the dynamics within this hierarchy, transitions between wells with relatively low barriers will occur so many times during one measurement that they will effectively average into a single state and not be observed in the experiment. In this manner, experimental details effectively determine the level of mechanistic detail that can be inferred from time-dependent, single-molecule data. Thus, the experimental distinguishability of the underlying molecular mechanism (*i.e*., the number of wells that can be separately observed in the experiment), as scaled by the experimental specifics of the measurement (see below), generates additional ‘observational’ DoFs for the experimental molecular mechanism.

### Hidden Markov models approximate the kinetic behavior of a single molecule

Having described how a molecular mechanism can be mapped onto a conformational free-energy landscape, we now discuss the process of inferring the details of such mechanisms from single-molecule signal *vs*. time trajectories. This inference requires that we approximate the time-dependent changes of the observed signal using some kinetic model, such as an HMM. HMMs are probabilistic models that describe a dynamic phenomenon that cannot be directly observed and is thus ‘hidden’ in the data (6). For our purposes, the hidden processes are the conformational dynamics of the biomolecules that are indirectly reported on by single-molecule signal *vs*. time trajectories. In an HMM, every data point in the signal *vs*. time trajectory corresponds to a particular ‘hidden’ state (*e.g*., a conformational state). However, the identities of these hidden states are unknown as the noisy signal is simply a proxy for the hidden states. For each of the possible hidden states, the HMM uses an ‘emission’ probability distribution to describe how that hidden state, as dictated by the experiment (see above), should appear in the noisy, observed signal (Figs. 2a and S2). The signal *vs*. time trajectory can then be approximated as a time-dependent sequence of hidden states (*i.e*., a Markov chain). In a Markov chain, transitions between the states, or between the same state (*i.e*., self-transitions), are stochastic and occur at random times. An HMM describes the probabilities of each of these transitions between the hidden states by assuming that the ‘transition probabilities’ are time-independent and depend only on the identities of the initial and final states (*i.e*., they represent a Markovian process exhibiting first-order kinetics (37)). These transition probabilities correspond to, and may be directly converted to, the respective transition rate constants for these processes (38). Thus, the general approach of an HMM aligns well with our above understanding of molecular mechanisms. Indeed, by separating transition and emission parameters, HMMs provide a framework wherein both the dynamics of the biomolecule (*i.e*., the mechanistic DoFs) and the experimental process by which these dynamics are observed (*i.e*., the observational DoFs) may be independently modeled and mapped onto one another.

To explore this equivalence between molecular mechanisms and HMMs further, we consider the hypothetical biomolecule described above that transitions between an open and closed conformation. In a single-molecule experiment, this system will yield a signal for each state (*e.g*., 0 for open and 1 for closed) and the resulting signal *vs*. time trajectory generated by such a molecule (Fig. 1a) may be described by an HMM (38, 39) (Fig. 2). Specifically, the emission probability distributions, which are often Gaussian distributions with mean *μ*and standard deviation *σ*, describe how the conformational states of the biomolecule manifest in the observed, experimental signal. Similarly, the rates of transition between the two conformations are described using transition probabilities between the two states, *P*_01_and *P*_10_, where *P*_*ij*_ is the transition probability from state *i* to state *j* between adjacent measurements in the signal *vs*. time trajectory.

HMMs have more parameters than the ones described above (Fig. 2a), but no more are required to describe this molecular system at equilibrium. The remaining parameters may be derived from constraints based on our prior knowledge of the experiment, and are thus not independent parameters. For our two-state example, the molecule can either remain in its current state (*i.e*., undergo a self-transition) or transition to the other state. These two options comprise an exhaustive set of mutually exclusive events. Thus, the probabilities of self-transition, *P*_00_ and *P*_11_, are

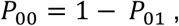

and

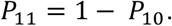

Similarly, for a system at chemical equilibrium, the probability that the molecule occupies either hidden state at the start of the experiment, π_0_ and π_1_, should be the steady-state probabilities

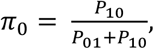

and

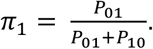

Therefore, while an HMM with two hidden states uses six parameters (*i.e*., the four *P*_*ij*_ and two π_*i*_) to model the observed dynamics, there are only two independent parameters if the system is at equilibrium (*i.e*., *P*_01_and *P*_10_); we call these the ‘transition’ DoFs. In general, an HMM with *K* hidden states will have *K*.(*K* − 1) transition DoFs (see Supplementary Information). Thus, a two-state HMM has two transition DoFs, which correctly matches the two mechanistic DoFs required to describe the conformational free energy landscape of our hypothetical two-state molecule (Fig. 1a).

We can similarly quantify the DoFs associated with the signal emitted from each hidden state in an HMM (*i.e*., the ‘emission’ DoFs). While the emission DoFs scale with the number of hidden states,*K*, the exact number depends on the distribution chosen to represent the hidden states; ideally, this distribution correctly encapsulates the experimental details of signal measurement (*e.g*., noise introduced by a detector). The number of emission DoFs in a *K* -state HMM is, thus, *mK*, where the proportionality factor, *m*, captures the dependence on the details of measurement. For single-molecule data analysis, the most standard emission distribution for a hidden state is a univariate (one-dimensional) Gaussian distribution. In this case, each hidden state is characterized by two emission DoFs (*m* = 2): one for the state mean, μ, and one for the state standard deviation, *σ*. In this work, we use univariate Gaussian distributions for the emissions of our hypothetical two-state biomolecule; emissions originating from the open state are modeled by a Gaussian distribution with parameters *μ*_0_ and *σ*_0_, and those from the closed state with *μ* _1_ and *σ*_1_ (Fig. 2a). Thus, our two-state HMM has four total emission DoFs, which correctly matches the four observational DoFs required to describe a typical detector-based measurement of two experimentally distinguishable states (Fig. 1a). However, emissions can be modeled by other sorts of distributions (40). Indeed, previous work on single-molecule fluorescence resonance energy transfer (smFRET) data analysis saw more accurate results by using a multivariate (two-dimensional) Gaussian emissions model for the fluorescence intensities of the donor and acceptor fluorophores (*I*_*D*_ and *I*_*A*_, respectively) *vs*. a univariate Gaussian emission model for the normalized FRET efficiency 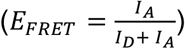, because of the flexibility provided by the additional emission DoFs.

In our example above, we have discussed using a two-state HMM with four emission DoFs and two transition DoFs to model a two-state molecular mechanism. However, one can always employ more complex kinetic models with more emission and transition DoFs to explain the dynamics of a biomolecule (Figs. S2 and S3, and Supplementary Information). In the ideal scenario, the transition DoFs of the HMM used to analyze a single-molecule experiment should match the corresponding mechanistic DoFs of the molecular mechanism under investigation, while the emission DoFs should match the observational DoFs of the experiment. As described in the next section, however, experimental complications frequently result in mismatches between the transition DoFs and mechanistic DoFs and/or between the emission DoFs and observational DoFs.

### Heterogeneity in a single-molecule ensemble reduces the apparent observational DoFs

In the previous section, we discussed how the transition and emission DoFs of an HMM may exactly match those of the mechanism underlying the dynamics of an individual molecule. However, this is only true in ideal cases in which each conformational state of the molecule gives rise to a unique, distinguishable signal state. In many experimental scenarios, this is not the case. In particular, heterogeneity within a biomolecular system can significantly complicate this process of kinetic modeling. Molecular heterogeneity is a well-documented phenomenon that affects biomolecular dynamics (33) in ways that are observable using many types of single-molecule techniques (31, 39).

While there exist many possible sources of heterogeneity, arising either from relevant changes to the biomolecular mechanism itself or due to experimental modalities such as surface tethering, we define this phenomenon explicitly as any experimental circumstances which cause a change in the free-energy barrier(s) between conformational states and, therefore, in the rates of transitions between them. While this is a somewhat restrictive definition, it covers a large number of situations that are observed in single-molecule studies (28–30) and allows us to quantitatively describe the effects of heterogeneity on the mechanistic and observational DoFs used to describe a biomolecular system. Specifically, we will use this definition to discuss two types of heterogeneity in the section below.

We first consider the scenario where changes to the free-energy barriers are time-independent—a condition referred to as ‘static heterogeneity.’ For our hypothetical biomolecule undergoing open⇌closed transitions, this type of heterogeneity could occur if a fraction of the biomolecules has undergone an effectively irreversible chemical change, such as a post-transcriptional or post-translational modification, or even some form of chemical damage. As a result of this change, the affected biomolecules may still undergo open⇌closed transitions, but they do so at a different rate than the unaffected molecules. In this case, the entire collection of molecules probed in the experiment (*i.e*., the experimental ensemble) consists of two subpopulations that undergo the same structural rearrangement, but at different rates—one slow and one fast (Fig. 1b). The conformational free-energy landscape of the molecules in this experimental ensemble can be understood as having split into two regions, each with their corresponding open- and closed-state wells, that are separated by a nearly infinitely high free-energy barrier across which transitions are not allowed on the experimental timescale. Thus, both of these subpopulations are separate two-well systems that are each characterized by two mechanistic DoFs (see above). Since the two subpopulations do not exchange, one additional mechanistic DoF is required to describe the fraction of each of the two subpopulations that comprise the ensemble. To match this mechanism, the corresponding kinetic model therefore requires five transition DoFs.

An additional complication arises when we consider that the chemical change causing the molecular heterogeneity might only alter the dynamics and not be drastic enough to alter the signal for each state in the two subpopulations. An HMM which fully captures this four-state system should, in the case of univariate Gaussian emissions (with *m* = 2), have eight total emission parameters— the four state means 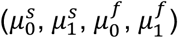 and the four corresponding state standard deviations 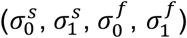 where the superscript stands for <u>s</u>low- or fast-transitioning molecules. However, due to the lack of differences between the signals for the states of the two subpopulations, *i.e*., between 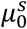 and 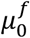, and 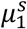 and 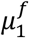 (and 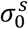 and 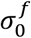, and 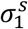 and 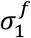), the number of apparent hidden states is reduced from four to two. The static heterogeneity in this case, therefore, leads to a mismatch between the apparent number of observational DoFs based on the observed signal states in the experimental dataset (four DoFs) and the expected number of observational DoFs based on the number of states in the underlying mechanism (eight DoFs) (Fig. 1b).

A similar outcome is also seen in the case of ‘dynamic heterogeneity,’ where the changes in the free-energy barriers are time-dependent. Our hypothetical open⇌closed biomolecular system could exhibit dynamic heterogeneity if the process that creates the slow and fast subpopulations is a reversible change, such as a slow orthogonal conformational rearrangement or the binding of a secondary factor that allosterically modulates the open⇌closed transitions. Such dynamic heterogeneity can cause molecules of the experimental ensemble to transition between a slow and fast phase of the open⇌closed rearrangements (Fig. 1c). The corresponding conformational free-energy landscape has four wells distributed along two reaction coordinates—one for the open⇌closed transitions and the other for the orthogonal slow⇌fast phase transitions. For our hypothetical biomolecule, we have chosen that the free-energy barriers separating the slow and fast phases are higher than those separating the open and closed conformations in each phase. This causes the transition rates between the slow and fast phases to be smaller than those between the open and closed conformations and creates a hierarchical separation of timescale between the two molecular processes (20, 33, 35). The dynamics of each of the two phases are described by two mechanistic DoFs (see above). Unlike the case of static heterogeneity, however, two additional mechanistic DoFs are used to describe the transitions between the two phases. To match this mechanism, the appropriate kinetic model, therefore, requires six transition DoFs.

A commonly occurring complication for the case of dynamic heterogeneity is that many single-molecule experiments are designed to produce signal changes along only one of the reaction coordinates. As a result, orthogonal processes described by the other reaction coordinate(s) may not lead to a change in the observed signal. Just as we described for static heterogeneity, the expected number of emission DoFs for an HMM with univariate Gaussian emissions (with *m* = 2) that describes this four-well system should be eight. However, if the transition between the slow and fast phases is not directly observable in the signal *vs*. time trajectories, just like the case for static heterogeneity, the apparent number of hidden states in the experimental dataset is reduced to two. This, therefore, causes a mismatch between the apparent number of observational DoFs based on the observed signal states in the experimental dataset (four DoFs) and the expected number of observational DoFs based on the number of states in the underlying mechanism (eight DoFs) (Fig. 1c).

Static and dynamic heterogeneities both cause discrepancies between the apparent number of observational DoFs based on the observed signal states and the mechanistically expected number of observational DoFs for a single-molecule signal *vs*. time dataset. These discrepancies obscure the underlying molecular mechanism and interfere with the process of kinetic modeling. For instance, in our examples of heterogeneity given above, a two-state HMM would match the apparent number of observational DoFs in the dataset, but would not have enough transition DoFs to match the molecular mechanism in the case of either static or dynamic heterogeneity. To be explicit, a two-state HMM has two transition DoFs, but the statically and dynamically heterogenous four-state systems that appear as two-state systems have five and six mechanistic DoFs, respectively. One approach to tackling such discrepancies is to select a model which has the correct number of transition DoFs (*i.e*., a four-state model in this case) and constrain the emission DoFs to match the apparent observational DoFs of the dataset. Hierarchical HMMs, which can be thought of as trees of HMMs (Fig. S3a), are a class of such models that have been used to analyze a diverse set of phenomena, including English language, cursive handwriting and musical pitch structure (41–43). Recently, in the field of single-molecule biophysics, they have successfully been employed to tackle the analysis of smFRET data containing dynamic heterogeneity (20). For hierarchical HMMs, the applied constraints reduce the emission DoFs in the kinetic model relative to a standard HMM with the same number of hidden states (see Supplementary Information) (44). Additionally, other physico-chemical constraints such as detailed balance can be applied to standard HMMs to address similar issues (45).

### Autocorrelation functions represent the dynamics of kinetic models

In this work, we investigate how different types of HMMs perform when faced with simulated single-molecule datasets where the states of the underlying molecular mechanisms have been obscured by different types of heterogeneity. The most straightforward manner in which this could be achieved is through direct comparisons between the distributions of HMM parameters inferred from simulated datasets and the simulated ‘true’ parameter values. Yet, for some of the HMMs that we investigated, the inferred kinetic model (for example, Fig. S2) was fundamentally different from the model used to generate the simulated dataset (Fig. S1). In these cases, a direct comparison between the model and ‘true’ parameters is not possible. As such, we sought to use a method capable of visualizing the dynamics specified by a kinetic model in a single analytical form to enable direct comparisons.

Autocorrelation functions (ACFs) have long been used to capture the dynamics of a system in a model-agnostic manner (46). An ACF provides a formal mathematical description of a time-dependent signal (*e.g*., a single-molecule signal *vs*. time trajectory, or a kinetic model) by analyzing the fluctuations between all pairs of points within the signal that are separated by a particular time difference (*i.e*., a lag time, τ). By performing this analysis as a function of the lag time, an ACF reports on the complete kinetic behavior of a signal. Fortunately, the ACF of an HMM can be calculated directly from the parameters of the HMM (see Supplementary Information), and so all of the information contained within an HMM can be represented in a single mathematical form—the ACF (Fig. 2b). Thus, we were able to represent the dynamics of the mismatched kinetic models that we investigated below by using their ACFs, which allowed us to compare the performance of kinetic models regardless of how the models were parametrized. While ACF-based analyses have been previously used for single-molecule experimental data (47), in this work, we have employed them primarily as a visualization and comparative tool to evaluate how well disparate kinetic models can capture the entire range of dynamic information contained within a single-molecule experimental dataset.

## Results and Discussion

### Development of an analysis platform that facilitates the application of multiple kinetic models

The analysis of single-molecule kinetics experiments requires several computationally challenging steps, of which, kinetic modeling usually represents the penultimate or ultimate step. Before any kinetic modeling may be performed, however, the raw data must be pre-processed and curated to eliminate spurious signal *vs*. time trajectories and generate a dataset that represents the ‘true’ mesoscopic ensemble (48–50). Subsequently, the data, the applied models, and the dynamics underlying the data and quantified by the models are visualized and evaluated. At present, there exist multiple computational pipelines that are capable of independently pre-processing, modeling, and visualizing single-molecule datasets (32, 51). These pipelines mostly implement their own, usually HMM-based, kinetic models. However, because these implementations have been developed independently, at different times, and by different research groups, they are not necessarily interoperable. This is also true for implementations of kinetic models that serve as independent platforms and are not part of specific pipelines. Even if they can be used interchangeably, switching between pipelines and platforms still serves as a barrier, since the outputs, in terms of the resulting model parameters and visualizations, are not always comparable. To generally facilitate the analysis of all single-molecule experiments and to specifically enable our investigation of the abilities of different HMMs to accurately infer the kinetics of ensembles exhibiting various types of heterogeneity, we have developed tMAVEN. As a flexible, open-source platform written in Python, tMAVEN can be employed for the processing, modeling, analysis, and visualization of single-molecule time-series data from a variety of experimental techniques and contexts.

tMAVEN offers capabilities for pre-processing and curating raw experimental data, in addition to generating multiple plots to visualize both the experimental data and the applied models. In this work, however, we focus on its ability to apply multiple kinetic models interchangeably from a single platform. While we have utilized several of the HMMs included in tMAVEN for our investigations here (12, 44), the architecture of tMAVEN is capable of handling any type of kinetic model which explains the dynamics of a mesoscopic ensemble of biomolecules as discrete transitions between relatively long-lived biomolecular ‘states.’ For instance, non-HMM-based kinetic models, such as thresholding and Gaussian mixture model-based clustering, are currently implemented as kinetic models in tMAVEN. In fact, because tMAVEN is an open-source and extensible platform, any analysis strategy that is congruent with this broad definition of a ‘kinetic model’ and that may be implemented or wrapped in Python code can be integrated into tMAVEN (see Supplementary Information). Crucially, we have standardized the outputs of the kinetic modeling functions in tMAVEN to yield a common set of parameters that describe the dataset as an ensemble of molecules (Fig. S4). Among other things, this standardization enables the presentation of any kinetic model with a common set of visualizations (*e.g*., population-weighted emissions distributions overlaid on a histogram of the data). Generally, these standardization requirements are just for the purpose of visualization or subsequent analysis steps (*e.g*., dwell-time distribution analyses), and they do not interfere with the inference or implementation of a kinetic model itself.

In this work, the standardization of kinetic model outputs in tMAVEN has also allowed us to easily collect the results of our kinetic modeling a variety of simulated datasets using different types of HMMs, and then to calculate distributions of the resulting kinetic model parameters (see Supplementary Information). Subsequently, comparing these parameter distributions and the inferred ACFs to the true parameter values and corresponding true ACFs enabled us to not only investigate the abilities of different types of HMMs to infer the underlying kinetics present in various heterogeneous ensembles, but also to highlight the regimes in which certain HMMs failed to accurately capture the kinetics of a particular ensemble.

### Global analysis allows accurate estimation of long-timescale dynamics from ensembles of short trajectories

We first evaluated the practice of combining signal *vs*. time trajectories into a single model (*e.g*., as in Refs. (24–27)) by investigating the analysis of simulated datasets of homogenous biomolecules (Fig. S1 and Supplementary Information). The simulated datasets were mesoscopic ensembles (*i.e*., composed of 100-1000s of identical molecules) that exhibited Markovian dynamics and had experimentally optimal signal-to-noise ratios and kinetic rates. Altogether, these properties represent the most ideal situation that one could expect to encounter when using an HMM to extract biomolecular dynamics from a single-molecule experimental dataset. Using these datasets, we compared two common methods by which an ensemble-level HMM is inferred which we call the ‘composite HMM’ and the ‘global HMM’ methods. In the composite HMM approach, an HMM is separately inferred for each individual signal *vs*. time trajectory, and the results from these individual HMMs are then composited together to generate an ensemble-level kinetic model (see Supplementary Information for details). In the global HMM approach, all individual trajectories are assumed to describe molecules undergoing dynamics corresponding to the same free-energy landscape and are thus independent and identically distributed according to the same underlying HMM (see Supplementary Information for details).

Surprisingly, we find that the composite HMM approach yields a result that is non-trivially different from the global HMM approach (Fig. 3a). The composite HMM appears to overestimate the transition probabilities for the mesoscopic ensemble, leading to faster decays in the mean inferred ACF when compared to the true ACF. This deviation is absent for the corresponding results from the global HMM analysis. While somewhat surprising, this result recapitulates the findings of previous investigations where a composite model was seen to be less accurate than a similar global model (40). We find that this overestimation of the transition probabilities is strongly correlated with the length of the trajectories comprising the single-molecule dataset (Fig. 3b and Fig. 4a). Interestingly, we also find that the overestimation is independent of the number of trajectories present in the dataset, with datasets containing fewer trajectories showing the same amount of deviation as datasets containing more trajectories (Fig. 3b and Fig. 4b). In both the composite and global approaches, the precision of the estimation depends on the amount of data, both in terms of lengths and number of trajectories. However, the accuracy of the global approach was notably independent of either length or number of trajectories (Fig. 3b and Fig. 4).

**Figure 3.**
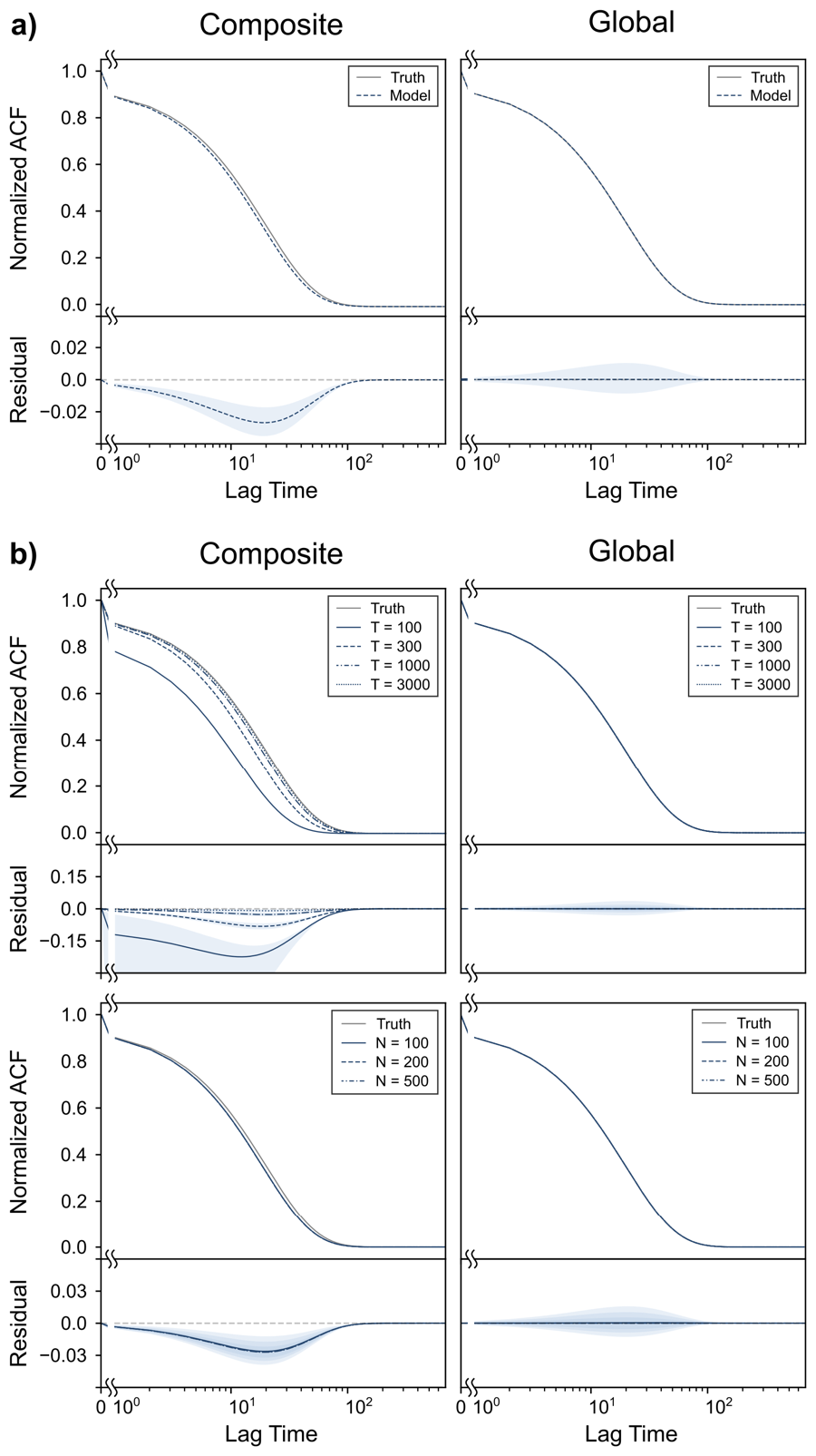
Comparisons of ACFs for homogeneous ensembles. **(a)** (top) The true ACF for the homogenous dataset (solid black) along with the mean of the ACFs (dashed blue) calculated using HMMs inferred from 10 ensembles using composite HMMs (left) and global HMMs (right), along with (bottom) the corresponding mean (dashed blue) of the residuals of the inferred ACFs to the true ACF. The blue area denotes the region one standard deviation away from the mean. The grey dashed line corresponds to zero. **(b)** The true (black) and model (blue) ACFs, along with the means of the residuals (blue), inferred using composite (left) and global (right) HMMs for homogeneous datasets of signal vs. time trajectories of varying lengths (top) and varying numbers (bottom). The blue area denotes the region one standard deviation away from the mean. The grey dashed line corresponds to zero.

**Figure 4.**
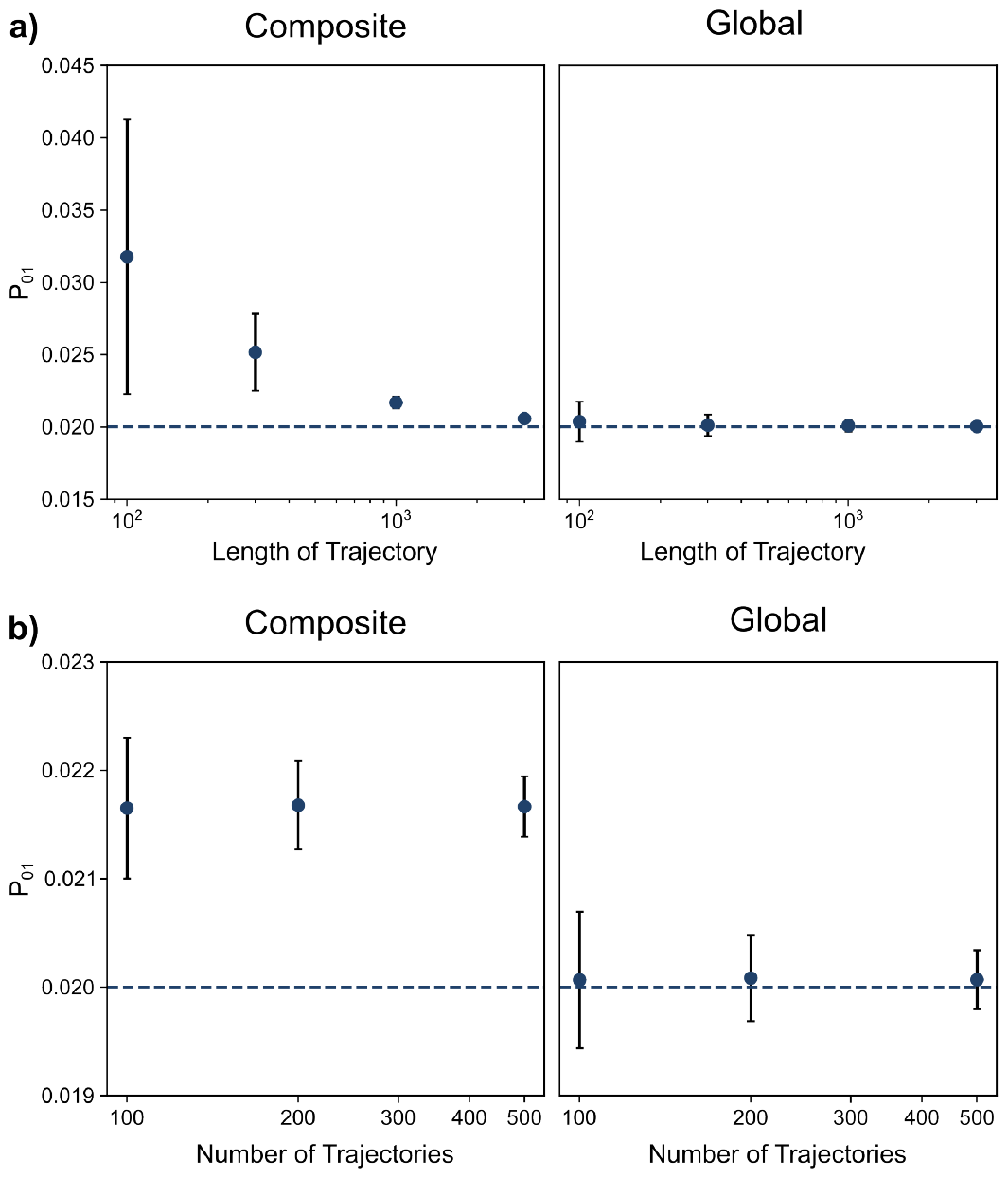
The effects of the lengths and number of trajectories in a mesoscopic ensemble on kinetic modeling. The transition probabilities from the ‘0’ state to the ‘1’ observed states inferred using (left) composite HMMs and (right) global HMMs from homogenous datasets with **(a)** varying lengths of trajectories and **(b)** varying numbers of trajectories. The dashed line represents the true transition probability for the dataset. The transition probabilities from the ‘1’ state to the ‘0’ state follow the same trend (data not shown).

This deviation in the accuracy of the composite HMM may be rationalized when we consider the effect of the length of the trajectories on the observed dynamics. In the case of HMM-based kinetic models, transition probability estimates are based on the apparent number of transitions that are observed between the states (see Theory and Supplementary Information). For very short trajectories, the sampling error in the number of observed transitions (both self-transitions and transitions to the other state) render our estimate of the underlying transition probabilities inaccurate for the composite HMMs. This is similar to the dwell-time distribution analysis scenario where ‘faster’ events (*i.e*., events with shorter dwell times) are over-represented in comparison to ‘slower’ events (*i.e*., events with longer dwell times) in short trajectories, which results in an over-estimation of the corresponding transition rates. We note here that the deviations are all over-estimations due to the regime of transition probabilities and trajectory lengths used for the simulated datasets. While those values correspond to commonly observed situations in single-molecule experiments, they also happen to fall in the regime where transitions between states are oversampled relative to self-transitions between the same state. Thus, this situation demonstrates a fundamental limitation to the amount of kinetic information present in a single trajectory.

To understand why this overestimation is not seen in the case of the global HMM, we analyzed the differences in the DoFs of these kinetic models. Both the composite and global HMMs have six total DoFs (two transition DoFs and four emission DoFs). However, for the composite HMM, individual HMMs are inferred for each trajectory before being composited into a single model. During this first inference step, the composite HMM uses 6N total DoFs, where N corresponds to the number of trajectories. In this step, there may or may not be sufficient information to accurately infer all 6N DoFs, especially when the individual signal *vs*. time trajectories are short. Regardless of how accurately they have been inferred, these 6N DoFs are reduced to six total DoFs in the subsequent compositing step and one should expect that any inaccuracies in the individual HMMs are propagated into the composited kinetic model. On the other hand, the global HMM consists of just six total DoFs throughout, and thus simultaneously integrates information from the entire ensemble to infer the corresponding kinetic model. While a short individual trajectory may not appear to be in the correct steady state, a global HMM accesses the steady state represented within the entire ensemble of molecules, and thus can accurately infer the underlying kinetics.

These results suggest that increasing the number of signal *vs*. time trajectories in a dataset is only advantageous if a global HMM approach is employed to simultaneously incorporate information from multiple trajectories into the analysis. Furthermore, these results also suggest that the correct *K*-state global model can accurately infer the kinetics of the underlying molecular mechanism even from datasets composed of very short trajectories (Fig. 4). This ability comes from the fact that the constraints used in the global HMM (*i.e*., all 6N DoFs are the same six DoFs) are applied throughout the entire inference process, unlike the composite HMM, in which the constraints are applied in a secondary step following the initial inference of the kinetic models. Interestingly, these constraints are equivalent to assuming that the ensemble under investigation is ergodic (*i.e*., all states of the underlying conformational free-energy landscape are accessible in the experimental timescale) (Fig. 1a). Thus, multiple short trajectories can be analyzed together, as though they were all generated from the long-timescale behavior of a single molecule. Such ergodic constraints, while key to the process of kinetic modeling, arise solely from our prior understanding of the molecular mechanism under investigation. The use of ergodicity as a constraint demonstrates how mechanism-informed modeling can improve the accuracy with which the kinetics present in a single-molecule dataset may be inferred.

### Constraints on emission distributions are required to accurately characterize static heterogeneity

Having evaluated the abilities of HMM-based modeling strategies to accurately estimate the kinetics of a homogeneous mesoscopic ensemble, we next sought to evaluate the performances of these methods in the presence of varying amounts of static heterogeneity (Fig. 1b). For this purpose, we simulated datasets where each signal *vs*. time trajectory had the same signal characteristics, but a subpopulation of trajectories had ‘fast’ open⇌closed transitions and the remaining had ‘slow’ open⇌closed transitions (Fig. S1). In this case, each trajectory is individually Markovian, because the transition probability depends only on the hidden state of the molecule (*i.e*., open or closed). However, since there are two subpopulations of molecules and thus two variants of each hidden state (*i.e*., slow or fast), the transition probability is not globally constant; it does not depend solely on the hidden state, rather it also depends on the subpopulation that the molecule belongs to. Furthermore, as is clear from the conformational free-energy landscape (Fig. 1b), all of the states in this mechanism are not accessible due to the presence of the near-infinite barrier separating the two subpopulations (see Theory). Therefore, the ensemble taken as a whole violates both Markovian and ergodic assumptions.

The effects of violating these assumptions can be clearly seen when we use a two-state HMM to analyze the dynamics of these simulated, statically heterogeneous ensembles. Since each individual trajectory within the ensemble is Markovian and has two distinct signal states, a two-state HMM (with four emission DoFs and two transition DoFs) is perfectly parameterized to exactly model each trajectory. Indeed, the individual trajectory-level HMMs that feed into the composite HMM do accurately capture the dynamics of both the slow- and fast-transitioning single molecules (Fig. 5). We further see that the distributions of the transition probabilities for these trajectory-level models clearly separate into two populations that scale linearly with the true proportions of the subpopulations. This signifies that information regarding the statically heterogeneous subpopulations may be captured at the trajectory level. However, we see that the ability to differentiate these subpopulations is completely lost for two-state models at the ensemble level (Fig. 5). Neither the composite HMM constructed from the individual trajectory-level HMMs (Fig. S5) nor the global HMM (Fig. 5) has enough complexity (*i.e*., enough transition DoFs) to describe the kinetics of the two subpopulations. However, it is somewhat surprising that the ensemble behaviors of the composite and global HMMs are very similar. In both cases, the inferred transition probabilities linearly scale with the subpopulation averaged transition probability for a specific state. Given that the individual HMMs themselves contain information on the heterogeneous subpopulations, this shows that both the composite and global two-state HMMs lose this information about the underlying heterogeneity due to ensemble averaging.

**Figure 5.**
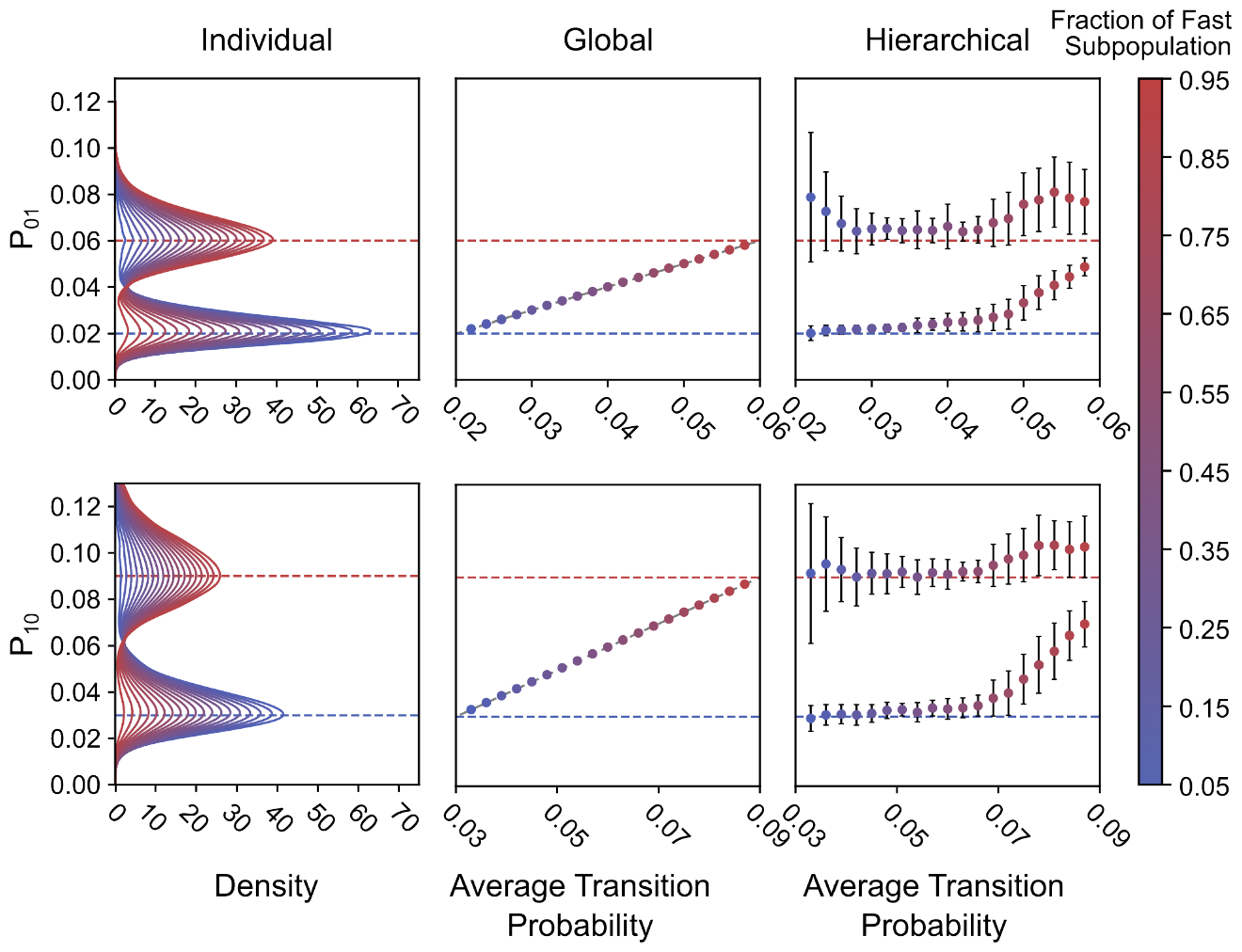
The effects of static heterogeneity on kinetic modeling. (left) Kernel density estimated distributions of the transition probabilities for the observed ‘open’ and ‘closed’ states inferred from the individual trajectory-level HMMs for each molecule in mesoscopic ensembles with varying amounts of static heterogeneity. Dashed red and blue lines denote the transition probabilities from each state for the subpopulation of fast- and slow-transitioning molecules respectively. (middle) The ensemble-level transition probabilities for the observed states inferred using global HMMs as a function of the average transition probability of the observed states (calculated using the proportions of fast- and slow-transitioning molecules). The dashed grey line denotes identity. (right) The two transition probabilities for each observed state as inferred using a hierarchical HMM as a function of the average transition probability of the observed states calculated using the proportions of fast and slow subpopulations.

Interestingly, both the four-state composite and global HMMs, which have the same number of hidden states as observational DOFs of the molecular mechanism (Figs. 1b and S2), are unable to differentiate between the two subpopulations. This failure of the four-state models lies both in the inference process and an intrinsic inability of this unconstrained kinetic model to capture the ensemble static heterogeneity. Without constraints to match the apparent four observational DoFs of the ensemble (see Theory), the four-state HMMs (with eight emission DoFs) do not computationally separate the observed emissions centered at 0.0 and 1.0 into the correct underlying hidden states (Fig. S6). Despite the flexibility provided by the additional DoFs, the inferred dynamics for the four-state HMMs are nearly identical to those for the two-state HMMs, when represented using their corresponding ACFs (Fig. S7). To further investigate this behavior, we initialized the inference process of the four-state global HMMs at the correct emission parameters (*μ*_*i*_ s and σ _*i*_s) and the correct steady state fractions (π _*i*_s). While this leads to a better estimation of the emission distribution themselves, where two emission means are clustered around 0.0 and two around 1.0 (Fig. S8a), the corresponding dynamics in the ACF remain unchanged (Fig. S8b). The major error is the inability to correctly identify and assign the observed datapoints to the slow or fast population, and this leads to a single, average Markovian behavior being assigned to states with the same emission means (Fig. S8c). Notably, this error persists even if the inference is initialized at all of the correct parameter values, including the correct transition matrix (Fig. S8d-f). Thus, further DoF constraints are required for accurate kinetic modeling.

Indeed, while the four-state models fail to correctly separate the observed emissions without fore-knowledge of the true values, the four-state hierarchical HMM (Fig. S3), which constrains the emission DoFs, is able to do so (Figs. 5 and S6). As a result, the four-state hierarchical HMM is the only ensemble kinetic model we used here that is capable of accurately capturing the statically heterogeneous subpopulations in the ensemble of trajectories. In particular, we see that it is most accurate for intermediate amounts of static heterogeneity where the fraction of fast subpopulation molecules in the ensemble is between 20 % and 60 %. Deviations from the true values of the transition probabilities outside of this regime can be explained by the hierarchical HMM assigning a non-zero probability to the rate of slow-to-fast interconversion at these levels of static heterogeneity. For our simulated ensembles exhibiting static heterogeneity, these probabilities should be zero, because the subpopulations do not interchange. However, since a constraint limiting this exchange is not explicitly applied for this hierarchical HMM, we observe that non-zero transition probabilities are inferred between the slow and fast subpopulations. Notably, this is the case even in the regime where the hierarchical HMM is the most accurate (Figs. 5 and S9). Instead of a near-infinite free-energy barrier separating the two subpopulations (Fig. 1b), the hierarchical HMM thus infers a finite barrier that may be traversed.

Altogether, these results highlight how essential it is to match the correct number of mechanistic and observational DoFs in a molecular mechanism when performing kinetic modeling. Here, static heterogeneity resulted in a reduction of the apparent number of hidden states from four to two (see Theory). A full, four-state HMM with eight emission DoFs was unable to compensate for this reduction. Being over-parameterized for the task, the four-state HMM was unable to correctly identify the two distinct emission states in the dataset. We found that only models with the correct number of emission DoFs were able to identify these states (*i.e*., both the composite and global two-state HMMs, and the four-state hierarchical HMM). Additionally, none of the HMMs we used had the same number of transition DoFs as the underlying molecular mechanism (*i.e*., five). The under-parametrized, two-state HMM with two transition DoFs yielded the correct ensemble-averaged transition probabilities but was unable to distinguish between the two subpopulations. On the other hand, the over-parametrized four-state hierarchical HMM with twelve transition DoFs (see Supplementary Information) was able to distinguish between the two subpopulations. However, the seven additional transition DoFs, which correspond to transitions between the two subpopulations, led to inaccuracies when one of the subpopulations had a significantly greater fraction than the other.

While beyond the scope of this work, our results suggest a simple approach to developing a kinetic model that exactly matches the transition DoFs of the molecular mechanism in the case of static heterogeneity. Since the heterogeneous subpopulations are captured in the individual HMMs with the matching emission DoFs (Fig. 5), a clustering algorithm can be used to classify the transition probabilities of these individual HMMs into a certain number of subpopulations (52), with the proportions of each cluster providing the additional transition DoFs; this is similar to, and could be done in conjunction with, the emission means-clustering approach we have used for the composite HMM. These results thus show how existing models like hierarchical HMMs (20) can be used to apply constraints that yield more accurate mathematical descriptions of static heterogeneity, while also serving as a guide for developing newer and better-performing kinetic models.

### Kinetic models can accurately describe dynamic heterogeneity when underlying processes are separated across timescales

Finally, we evaluated the abilities of HMM-based models to infer the kinetics of a mesoscopic ensemble exhibiting different levels of dynamic heterogeneity (Fig. 1c). For this purpose, we simulated datasets of trajectories in which molecules can interchange between a fast-transitioning and a slow-transitioning phase, both of which have the same emission properties (Fig. S1). Unlike in the case of static heterogeneity, each individual trajectory here appears to be non-Markovian. This is because, for each signal state (*i.e*., the set of hidden states with the same emission distributions), the transition probabilities depend on whether the molecule is in the fast-transitioning phase or the slow-transitioning phase (Fig. 1c). Furthermore, the ensemble, in this case, is ergodic since each molecule can access all of the wells in the underlying conformational free-energy landscape.

The differences between static and dynamic heterogeneity become clear when we consider how an individual, two-state HMM models the trajectories containing dynamic heterogeneity before being used to create the two-state composite HMM (Fig. 6). As the total rate of interchange between the two kinetic phases (defined as *P*_*sf*_ + *P*_*fs*_, where the subscripts stand for the slow and fast-transitioning phases) was increased, we found that the distributions of the individual transition probabilities between the open and closed states approached the ensemble average values for the slow and fast kinetic phases (Fig. 6). This asymptotic behavior highlights how the simulated trajectories more rapidly relax to the equilibrium behavior of the ensemble at higher rates of interchange. At lower rates of interchange, there are insufficient numbers of transitions between the two phases on the timescale of the experiment. As such, the individual trajectories, and thus the individual HMMs, more closely resemble the static heterogeneity case (Fig. 5). Indeed, this observation reflects how static heterogeneity can be thought of as a limiting case of dynamic heterogeneity where the transitions between the different kinetic phases occur much slower than the experimental timescale, making the free-energy barrier between the two effectively infinite within the experimental context (see Theory). Interestingly, we did not see an effect from varying the rates of interchange on the results of either the composite HMM (Fig. S10) or the global HMM (Fig. 6), because the dynamical effects caused by the underlying dynamic heterogeneity are not able to be captured by either of these kinetic models.

**Figure 6.**
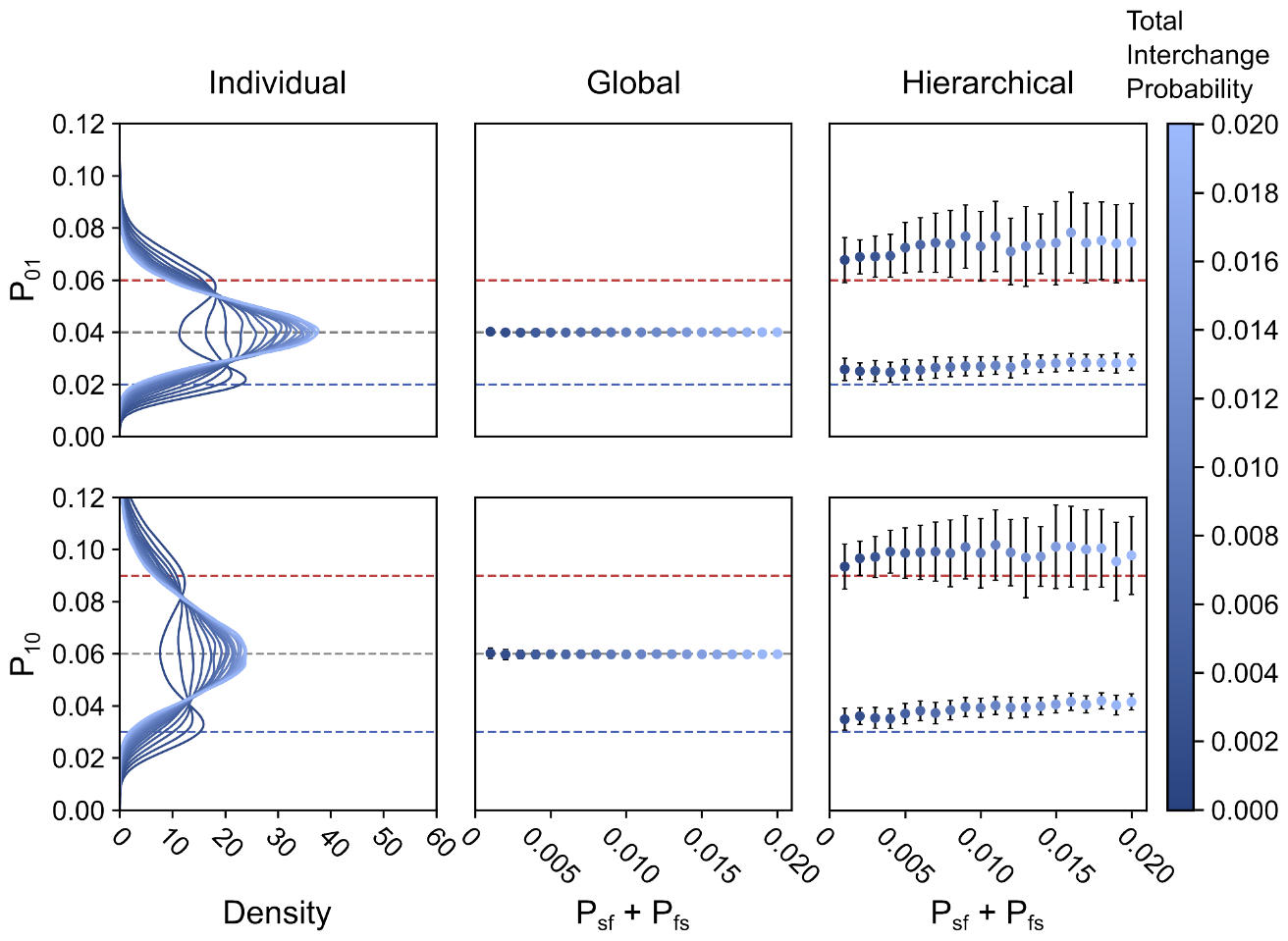
The effects of dynamic heterogeneity on kinetic modeling. (left) Kernel density estimated distributions of the transition probabilities for the observed ‘open’ and ‘closed’ states inferred by the individual trajectory-level HMMs for each molecule in mesoscopic ensembles with varying total probability of transition between slow- and fast-transitioning phases (P_sf_ + P_fs_). Dashed red and blue lines denote the transition probabilities of each state for the fast- and slow-transitioning phases respectively. The dashed grey line denotes the ensemble average transition probability of each observed state. (middle) The ensemble-level transition probabilities for the observed states inferred using global HMMs as a function of the total probability of transition between slow- and fast-transitioning phases. (right) The two transition probabilities for each observed state inferred using hierarchical HMMs as a function of the total probability of transition between slow- and fast-transitioning phases.

As we saw in the case of static heterogeneity, analyzing simulated datasets exhibiting dynamic heterogeneity using full four-state HMMs that lacked any constraints on the emission distributions yielded the same ACFs as the analysis using two-state HMMs (Fig. S11). This was caused by the same mischaracterization of the emission distributions of the hidden states as was observed for the static case (Fig. S12). Even initializing the inference process at the correct underlying model parameters was unable to address this error (Fig. S13). In contrast, we saw that analyzing the datasets with a four-state hierarchical HMM was much more successful and could accurately distinguish between the two kinetic phases (Figs. 6 and S12). Although, we note that increasing the rates of interchange did slightly decrease the accuracy of the transition probabilities between the observed signal states (Fig. 6). This inaccuracy appears to correlate with the inaccuracy in estimating the correct values of *P*_*sf*_ and *P*_*fs*_ themselves at higher rates of interchange (Fig. S14). Indeed, we saw that at lower rates of interchange, the inferred values of *P*_*sf*_ and *P*_*fs*_ themselves matched the simulated values, albeit with a slight systematic over-estimation, and they plateaued for fast interchange between the kinetic phases. We hypothesize that this is because at higher rates of interchange between the kinetic phases, the underlying molecular mechanism of the dynamic heterogeneity stops reflecting a hierarchical set of dynamics (Figs. 1c and S3a), and more closely resembles a system where transitions between all four states are equally temporally separated on the conformational free-energy landscape. Furthermore, the hierarchical HMM was able to model the statically heterogeneous ensembles with equal proportions of slow- and fast-transitioning molecules (Fig. 5) more accurately than the dynamic heterogeneous ensembles—which also have an equal average proportion of slow- and fast-transitioning molecules. Taking this statically heterogeneous dataset to represent the limit of infinitely slow interchange for the dynamic heterogeneity we have simulated here, our results thus suggest that a large separation of timescales between the directly observed molecular process (*i.e*., the open⇌closed dynamics) and the ones responsible for causing the dynamic heterogeneity (*i.e*., the slow⇌fast interchanges) is required for the hierarchical HMM to accurately infer kinetics from these datasets.

Once again, these results demonstrate that the number of transition and emission DoFs serves as a good rule of thumb to determine how different HMMs will perform at inferring the underlying molecular mechanism of a biological system. When analyzing single-molecule datasets, thorough consideration must be given to how the HMM DoFs correspond to the specific mechanistic and observational DoFs of the molecular mechanism(s) of interest. For example, while ensembles exhibiting static heterogeneity and dynamic heterogeneity differ by a single transition DoF (five and six, respectively), considering the apparent ergodicity of the resulting signals means that we should expect that each individual trajectory is also characterized by two or six transition DoFs, respectively. An individual two-state HMM matches the transition DoFs in the individual trajectories in the static case. However, two-state HMMs are under-parameterized while four-state HMMs, including hierarchical HMMs, are over-parameterized in the dynamic case. Thus, the only general approach to accurately extract heterogeneous kinetics and molecular mechanisms from single-molecule datasets using HMMs is to customize the HMM being used by applying constraints that match the transition DoFs and emission DoFs to the mechanistic and observational DoFs of the underlying molecular mechanism. Such customization requires that a researcher know whether they are expecting static or dynamic heterogeneity *a priori*, so we do not expect a universally applicable method to be easily developed. Regardless of these difficulties, approaching kinetic modeling of single-molecule datasets in this manner will greatly expand the richness of information that can be extracted from single-molecule experiments, as well as the types of biomolecular processes that may be studied using these techniques. Altogether, our results show that developing more accurate kinetic modeling strategies will require thoughtful approaches to applying physico-chemically informed constraints to presently existing models. For instance, the hierarchical HMM does apply constraints to the emission DoFs, but at present is still over-parametrized with twelve transition DoFs compared to the six transition DoFs in the dynamic heterogeneity molecular mechanism. Fully matching all of the mechanistic and observational DoFs of an experimental molecular mechanism will enable kinetic models to better replicate the underlying dynamics they seek to describe.

## Conclusions

In this work, we have shown how mismatches in the transition and emission DoFs of kinetic models and the mechanistic and observational DOFs of molecular mechanisms, respectively, play a role in determining the accuracy of modeling single-molecule dynamics. However, matching mechanistic and observational DoFs is only the first step in mechanism-informed modeling. True mechanism-informed modeling requires an in-depth formulation of how a model captures the underlying physico-chemical properties of both the biomolecular system of interest and the detection process that yields the experimental signal *vs*. time trajectory (53). For example, previous work has shown that accounting for the Poisson-distributed counting statistics of photon emission and the effects of detector noise affects the accuracy of kinetic modeling (40). Similarly, accounting for the integration time of a detector allows for the modeling of sub-temporal-resolution kinetics (54–59).

Taken together, a universal single-molecule kinetic model (32) is unlikely to be appropriate for all experimental contexts. Indeed, in this work, we have focused on analyzing relatively ideal, simulated datasets by using just two hidden states, Gaussian noise, optimal signal-to-noise ratio, a lack of temporally averaged measurements, a large number of long trajectories, and a clear mechanistic separation of static and dynamic heterogeneity. Yet, even under these ideal conditions, we quickly reached the limits of current HMM-based modeling approaches. Real experimental datasets are much more complex, with more hidden states, non-trivial noisy emission distributions, and kinetics that may be slower or faster than is experimentally accessible. Additionally, in many cases, the information in real datasets is limited in terms of the length (*e.g*., due to photobleaching) or number of trajectories (*e.g*., due to experimental throughput). In contrast to our simulated datasets, experimental datasets may contain both static and dynamic heterogeneity that simultaneously originate from different sources. In such cases, it is not immediately obvious what the underlying mechanism for a particular biomolecular system should be, which makes the use of solely HMMs to analyze such data nontrivial. In some of these scenarios, other analysis methods (*e.g*., dwell-time distribution analysis (60, 61)) may indicate the presence of heterogeneity and model-mechanism mismatch. However, in many cases, even if related biochemical experiments or alternative analysis methods indicate that a single-molecule dataset is heterogeneous, a mismatched model may remain the only way meaningful mechanistic information can be extracted from the experimental data.

In the absence of a universally optimal kinetic model, we propose that the most appropriate approach to kinetic modeling is to use multiple different kinetic models and compare the relative insight of their results to one another. Our investigation, which demonstrates how different types of models can be expected to perform in several cases of model-mechanism mismatch, can aid the interpretation of kinetic modeling results in situations where external evidence points to a model-mechanism mismatch, which may, in turn, guide the design of further experiments to avoid or minimize such mismatch. Of course, the seamless application and evaluation of multiple kinetic models to the same dataset is computationally and logistically challenging. This is especially true given that many kinetic models and their corresponding software implementations were developed at different times, in different research groups, and using different programming languages. The development and use of tMAVEN, which can interchangeably apply and visualize several different types of HMMs and other kinetic models, alleviates many of these challenges. At present, we have packaged tMAVEN with Python implementations of ∼15 kinetic models including upgraded versions of those used in previously published work (*e.g*., HaMMy, vbFRET, ebFRET, hFRET, *etc*.) (6, 11, 12, 16, 20) as well as in the current work. As an open-source, extensible program, we have also ensured that the core tMAVEN modeling functions are flexible enough to allow other modeling approaches to be easily added. We envision tMAVEN as a platform that can be used by the entire single-molecule biophysics community for the future development of single-molecule data analysis software, including the further development of some of the modeling approaches we discuss here. By allowing developers to make use of the pre-existing user interface and data processing functions in tMAVEN, we hope to streamline future advances in the kinetic modeling of single-molecule datasets. This accessibility, paired with tMAVEN’s user-friendly interface, lowers the barriers for both the use and development of kinetic modeling tools and thereby enables the type of mechanism-informed kinetic modeling that we envision will be so powerful for researchers using single-molecule experimental techniques to study biomolecular dynamics.

## Author Contributions

All authors designed the research, analyzed the data, and wrote the manuscript.

## Acknowledgements

We thank Chris Wiggins, Jake Hofman, Jonathan Bronson, Jan-Willem van de Meent, Jason Hon, Bridget Huang, and Riley Gentry for many insightful discussions and collaborations regarding the kinetic modeling of single-molecule experimental data and Jason Hon, Bridget Huang, Sukjin Jang, Riley Gentry, Nina Michael, Qiongfang Zhang, Robin Shivnaraine, John Janetzko, Jonathan Deutsch, and Qianyi Wu for feedback on the software features in tMAVEN. This work was supported by funds to R.L.G. from the National Institutes of Health (NIH) (R01 GM 084288, R01 GM 137608, R01 GM 128239, and R01 GM 136960) and the National Science Foundation (NSF) (CHE 2004016) as well as funds to C.D.K. from the NIH (Training Grant in Molecular Biophysics to Columbia University, T32 GM008281), the Department of Energy (DOE) (Office of Science Graduate Fellowship, DE-AC05-06OR23100), and the NSF (CHE 2137630).

## Code and data availability

The open-source code for tMAVEN is implemented in Python and is freely available *via* a Git repository accessible at https://github.com/GonzalezBiophysicsLab/tmaven. The Python code used to generate the simulated ensembles, interface with tMAVEN, and programmatically analyze the simulated ensembles using different HMMs as well as subsequently collect the inferred results and generate the plots published in this article is also freely available *via* a separate Git repository available at https://github.com/GonzalezBiophysicsLab/tmaven_paper. tMAVEN documentation and a user manual are at provided at https://gonzalezbiophysicslab.github.io/tmaven/, in addition to a video tutorial analyzing real-world, experimental smFRET data and sample scripts for various analysis

## Supplemental Information

### 1 Construction of simulated datasets

Many hidden Markov model-based (HMM-based) analysis approaches used for single-molecule kinetic analysis were developed specifically for single-molecule fluorescence resonance energy transfer (smFRET) experiments [1]. As such, we decided to generate simulated datasets with parameters that replicate an ideal time series that might be obtained from such smFRET experiments. Nonetheless, our datasets can also be thought of as a set of general normalized signal *vs*. time trajectories, and, therefore, our results remain applicable to datasets obtained using other single-molecule techniques (*e.g*., force-clamp spectroscopy, single-molecule field effect transistors, *etc*.).

For the dataset lacking heterogeneity (“Homogeneous”), we generated an ensemble of trajectories representing a hypothetical biomolecular reaction 0 ⇌ 1, where 0 and 1 are the respective labels for the open and closed states described in the main text (Figure S1). The emission distributions for this dataset (and all subsequent simulated datasets described here) were Gaussian distributions with emission means μ_0_ = 0.0 and μ_1_ = 1.0 for the states 0 and 1, respectively. The emission noise for these Gaussian distributions was 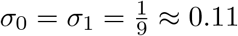. As a result, the signal-to-noise ratio (SNR) for these trajectories was 9. The steady-state probabilities were chosen to asymmetrically favor state 0 over state 1 with π_0_= 0.6 and π_1_= 0.4, resulting in K_eq_ ≈ 0.66. We chose to simulate these trajectories in a ’slow’ kinetic regime that replicated the experimental scenarios where the effects of time averaging, which negatively affect the effectiveness of single-molecule experiments [2, 3], would not be significantly present. Based on this constraint and the pre-defined steady-state probabilities, transition probabilities (written as *P*_ij_ for transitions from the initial state i to the final state j) were chosen to be *P*_01_ = 0.02 and *P*_10_ = 0.03. Due to constraints on normalization, the remaining self-transition probabilities were *P*_00_ = 1 *- P*_01_ and *P*_11_ = 1 *-P*_10_. These transition probabilities are large enough that multiple transitions between states 0 and 1 would occur within the length of a typical smFRET time series (*e.g*., 1000 datapoints). To replicate an ideal experimental dataset, each simulated dataset consisted of 200 individual trajectories of 1000 time-points each (unless otherwise stated). To ensure that our results were statistically reliable and accounted for the uncertainties due to sampling errors, we generated and analyzed 100 dataset variations for each ensemble using the same parameters. Details about the parameters used for this simulated dataset are shown in Figure S1. This Homogeneous dataset is a good representation of an ideal single-molecule experiment and thus is appropriate for investigating the analysis of kinetic information using HMMs.

To investigate heterogeneous ensembles, we simulated datasets containing static heterogeneity or dynamic heterogeneity (Figure S1). For the ensembles with static heterogeneity (“Static”), we simulated datasets composed of two different subpopulations of trajectories. Both subpopulations were characterized by the same emission distribution as the Homogeneous datasets; however, one exhibited ‘slow’ kinetics while the other exhibited ‘fast’ kinetics. The slow subpopulation consisted of trajectories generated with identical kinetic parameters to the Homogeneous ensemble above. The fast subpopulation had identical steady-state probabilities as the slow subpopulation but had transition probabilities between 0 and 1 states which are three times larger than the ones for the slow subpopulation (*i.e*., *P*_01_ = 0.06 and *P*_10_ = 0.09). To test the effect of varying levels of static heterogeneity, we varied the fraction of the fast subpopulation in these datasets from 5% to 95 %. Details about the parameters used for this simulated Static dataset are shown in Figure S1.

For the ensembles with dynamic heterogeneity (“Dynamic”), we simulated trajectories with inter-converting ‘fast’ and ‘slow’ kinetic phases. The emission probabilities for these two phases were identical to the Homogeneous and Static datasets. The transition and steady-state probabilities of the fast kinetic phase were the same as those for the fast subpopulation of the Static dataset, and those for the slow phase corresponded to the slow subpopulation. The transition probabilities of interchange between the fast (*f*) and slow (*s*) phases were varied such that the steady-state proportion of the two phases remained constant at a 1:1 ratio, and were increased to test increasing dynamic heterogeneity. This yielded *i.e*., *P*_*sf*_ = *P*_*fs*_ that varied from 0.0005 to 0.01. Details about the parameters used for this simulated Dynamic dataset are shown in Figure S1.

### 2 Calculation of Degrees of Freedom of HMMs

The number of degrees of freedom (DoFs) of a model refers to the number of independent parameters required to completely define it. The number of DoFs can be computed by considering the total number of parameters in the model and the number of constraints that need to be applied to these parameters based on prior knowledge of the system. DoFs provide useful insight into the complexities of kinetic models and how they can match the mechanism of a biomolecular system. Below, we calculate the DoFs for a general HMM and then show these can be applied to the specific cases described in the main text.

#### 2.1 General Approach for calculating DoFs for a standard HMM

A standard HMM consists of the following sets of parameters: the transition matrix ***A***, the initial probabilities π, and, for an HMM with univariate (one-dimensional) Gaussian emission distributions, the sets of emission means ***μ*** and emission standard deviations ***a-***. For an arbitrary *K*-state HMM, ***A*** will contain *K*^2^ entries to account for the probabilities for all possible transitions between the *K* states. In the ***A***s described below, each element represents the probability of transition between the initial state as given by the row index and the final state as given the column index (*i.e*., *P*_*initial,final*_). Thus,

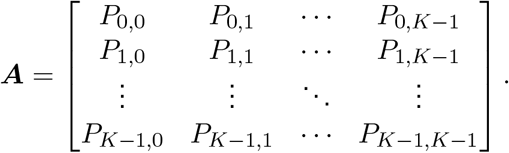

However, these elements in ***A*** are not all independent. Each row of ***A*** consists of the probabilities of all possible transitions from a particular state and thus must sum to one. By enforcing a row normalization constraint for each of the *K* rows, we are left with only the number of independent parameters required to define the transition matrix, which is seen to be *K*(*K -*1). In terms of molecular mechanism, this can be rationalized with the observation that transition rates are only defined between different states, and so, there is no rate constant (or free-energy barrier) for self-transitions. The corresponding self-transition probabilities (which are the elements along the diagonals of ***A***) can be thought of as consequences of the normalization of the transition probabilities to other states in the model. Thus,

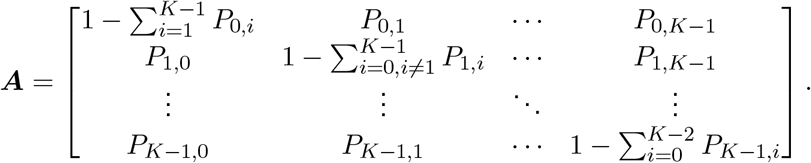

Similar to ***A*** above, a *K*-state HMM has *K* initial probabilities which can be represented by π. Yet, for a system at chemical equilibrium, it is immaterial when the observation starts, and thus each of the π_i_s (representing the initial probability of the *i*th state) is equal to the steady-state probability of the *i*th state, 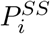, which in turn is wholly defined in terms of *A* by the relation

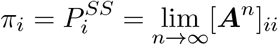

where the notation *B*_*ii*_ represents the element in the *i*th row and *i*th column of a given matrix ***B***. With these K constraints, therefore, we see that **π** can be expressed completely in terms of ***A***. Thus, while the kinetics of a *K*-state HMM has *K*^2^ + *K* parameters, they are also characterized by 2*K* constraints (*K* for ***A***, and *K* for **π**), leading to a net *K*(*K -*1) independent parameters (*i.e*., DoFs) that define the kinetics of these HMMs. We refer to these as the transition DoFs of the HMM.

Similar to the above, we can also determine the number of DoFs that characterize the emissions of an HMM, (*i.e*., the emission DoFs). While this choice scales with *K* for a given HMM, we see that the exact number of DoFs depends on the specific emission distribution used for the HMM [4], with a proportionality constant *m* that is characteristic of this emission distribution. For a univariate Gaussian distribution emission, each state *i* is characterized by a single emission mean μ_i_ and a single emission standard deviation σ_*i*_. These are typically independent for each state, and therefore ***μ*** and ***a-*** are together characterized by 2*K* DoFs (in this case, *m* = 2). It follows from this that HMMs which have other emission distributions would also have differing numbers of such emission DoFs. For example, HMMs with univariate Poisson (*m* = 1) or beta distributions (*m* = 2) for emissions have *K* and 2*K* emission DoFs, respectively. In the case of smFRET, where the individual trajectories may be analyzed as either two-dimensional independent intensities of a pair of donor and acceptor fluorophores (*I*_*D*_ and *I*_*A*_ respectively) or as the one-dimensional normalized FRET efficiency 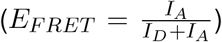, the two-dimensional case may also be represented by HMMs with a multivariate (in this case, bivariate) Gaussian emission distribution (*m* = 4). In this case, this HMM would be characterized by 4*K* emission DoFs. Crucially, while the exact number may vary depending on the specific emission distributions, for a standard HMM with no constraints on emission, the number of emission DoFs is proportional to *K*.

Based on the above framework, we can directly see, as mentioned in the main text (and Figure 2), that a two-state HMM with univariate Gaussian emission distributions will have 2 transition DoFs and 4 emission DoFs. Similarly, a four-state HMM with univariate Gaussian emissions (Figure S2) will have 12 transition DoFs and 8 emission DoFs. In the following section, we shall use the above approach to calculate the DoFs of a hierarchical HMM.

#### 2.2 Calculating DoFs for a two-level four-state hierarchical HMM

Hierarchical HMMs (hHMMs) are a category of kinetic models that can be thought of as HMMs within HMMs, across several levels. For this work and the ease of the following discussion, we shall focus in particular on a two-level, four-state hHMM (Figure S3a). The lowest level of the hHMM is called the production level (in this case, *d* = 3), and the states in this level are used to define the emission distributions of the hHMM. Since there are four states in this level, the entire hHMM is a four-state kinetic model. Crucially, however, unlike a four-state HMM (Figure S2), these four states are connected using two HMMs, which in turn are related using a higher level (*d* = 2) HMM. Because of this structure, in addition to the expected transition parameters for each of these HMMs (***A***s and **π**s), hHMMs also require a set of exit probabilities, ***E***s, that define transitions to a higher level. The root level (*d* = 1), in this case, functions like the initial node of an HMM (Figure 2 in the main text and Figure S2). Thus, for the two-level, four-state hHMM, the transition parameters can be arranged in families, based on their levels, as follows

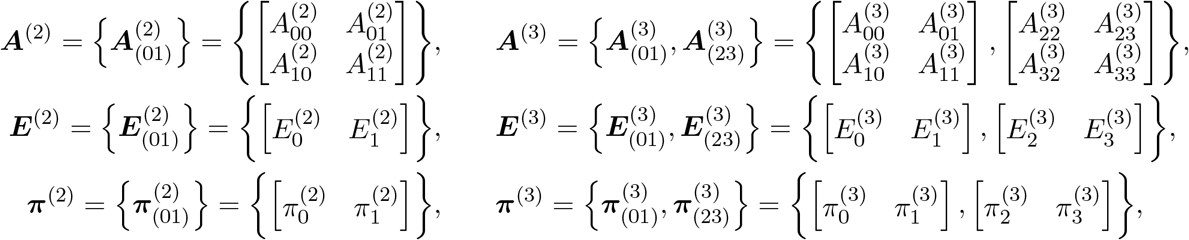

where the superscript (*n*) denotes the level of the hHMM and subscripts refer to the states of the HMM at that specific level (Figure S3a). Because of the hHMM structure, the parameters do not necessarily independently capture the probabilities of transition between states of the production level, as would be the case with a standard HMM. Instead, using these parameters, we can generate a four-by-four transition matrix (where each element is defined as above for the standard HMM) that describes the probability of all possible transitions between production states. It is important to note, that, in this case, for each transition between production states, multiple transition paths must be accounted for (Figure S3b). For example, the 0 state of the production level (*d* = 3) may self-transition in the production level itself 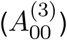, enter the higher level (*d* = 2), self-transition, and rejoin itself in the production level 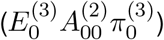 or journey vertically to the root node (*d* = 1) before returning to itself 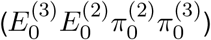. Thus, the total probability of self-transition for the production-level 0 state (*P*_00_) is the sum of these three paths. Repeating this process for all possible transitions between states in *d* = 3, we have:

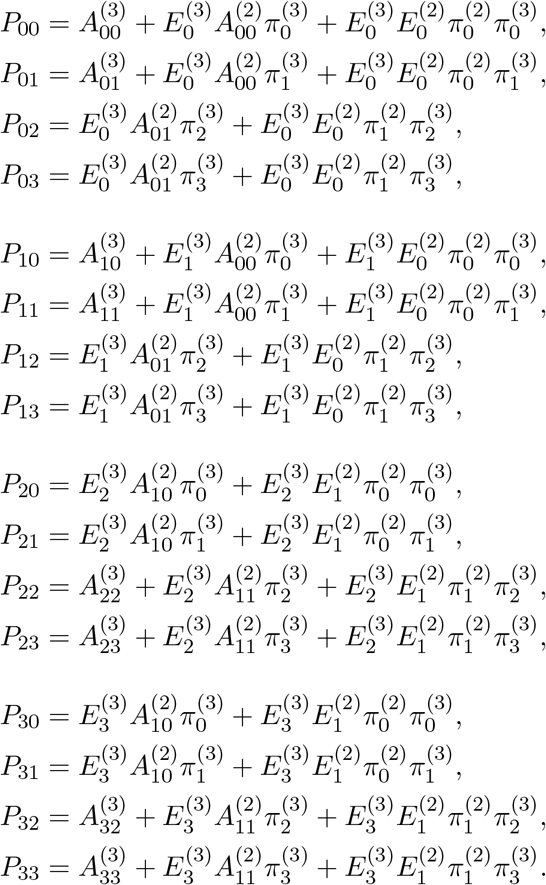

Based on the above, we see that two-state four-level hHMMs have, in total, 24 parameters that define the kinetics of these models. However, we also see that there are constraints on these parameters. For example, each HMM across all levels of the hHMM (three in this case) have normalization constraints [5]. First, the sum of the entry probabilities for all HMMs must equal one. In this case,

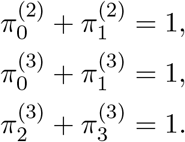

Second, the sum of all transitions and exits from a state at all levels must equal one. Thus

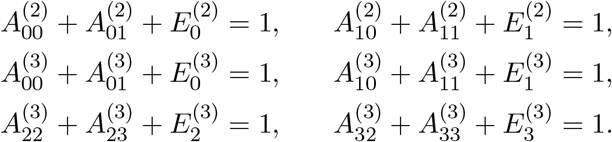

Imposing these nine normalization constraints on the corresponding transition probabilities, we get

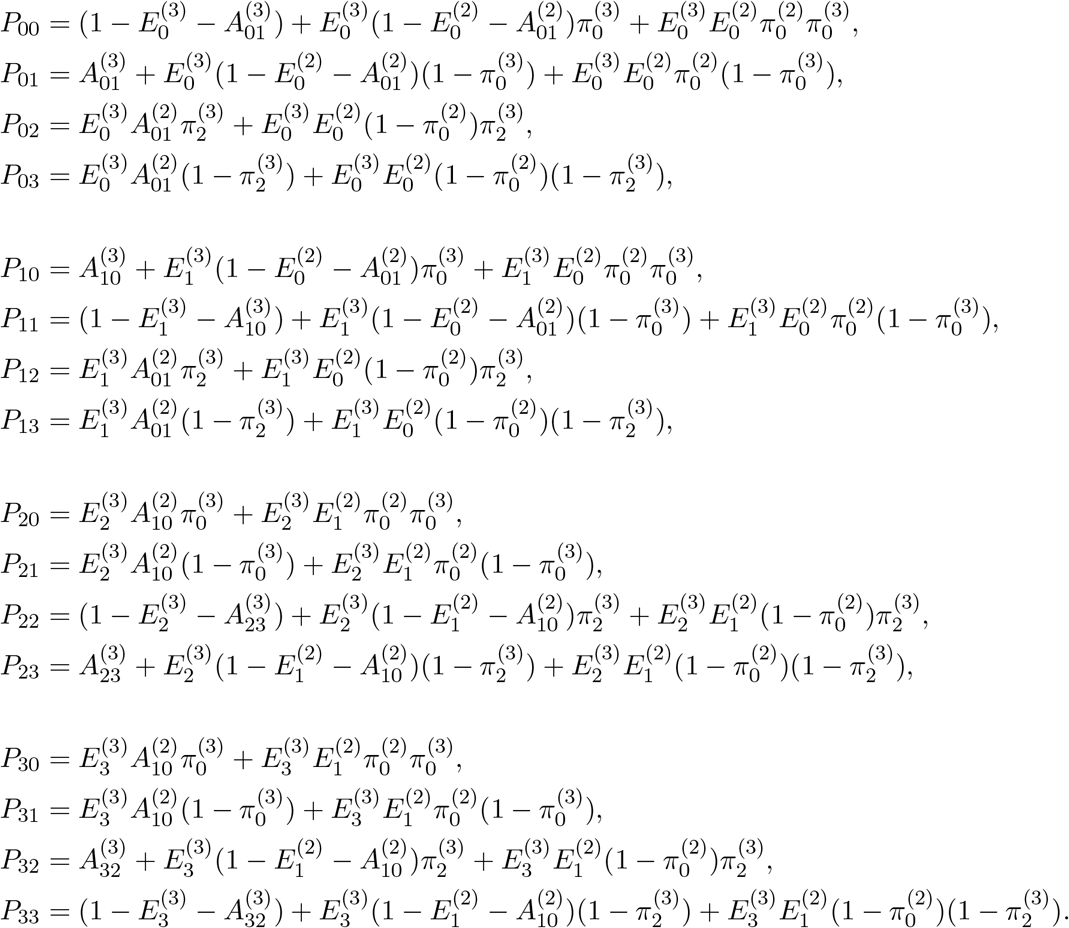

Apart from the above normalization constraints, we see that there are constraints on the steady-state probabilities of each of the HMMs across the different levels, which arise from the consideration that the underlying chemical system is in equilibrium. Unlike the standard HMMs however, the expressions for these steady-state probabilities are not straightforward, and require enumeration of the multiple paths that connect the states using the diagram method [6]. Nonetheless, these steady-state probabilities may be used to constrain one set of parameters (for example, the 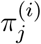 for state *j* in level *i*) with respect to the other parameters. Following the application of these constraints, we are left with an independent parameter count of 12: 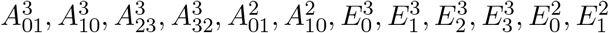. Thus, this hHMM has 12 transition DoFs.

The strength of hHMMs comes from the fact that these kinetic models may be used for inference in cases where the emissions between different production states are shared. In this case, even though the kinetic model possesses four underlying states, the emissions between states 0 and 2 are identical, as are the ones between states 1 and 3. Thus, this hHMM possesses only two univariate Gaussian emission distributions, resulting in 4 emission DoFs. This is what differentiates this two-level, four-state hHMM from a standard four-state HMM, which also possesses 12 transition DoFs but 8 emission DoFs. In the main text, we discuss the implications of this difference and how they result in the different abilities of these kinetic models to capture the underlying molecular mechanism of a single-molecule dataset.

### 3 Evaluation of ensemble dynamics using auto-correlation functions

One of the most straightforward ways to determine the accuracy of HMMs in modeling our simulated data is to compare the inferred HMM parameter distributions to the ‘ground truth’ HMM parameters used in the simulations. However, for several of our analyses, the HMM used during inference was fundamentally different from the generative model used in the simulation. The use of two-point auto-correlation functions (ACFs) enabled us to ascertain the ability of an HMM to quantitatively describe the dynamics of the entire ensemble of trajectories, even in the case of disparate underlying models.

Briefly, the signal *y* measured in a single-molecule experiment is a function of time (*y* = *y*(*t*)). A two-point ACF quantifies the correlation in the value of *y*(*t*) and the value of *y*(*t*+τ) after some lag time, τ, during which the signal evolves [7]. In its un-normalized form, this ACF is an expectation value given by

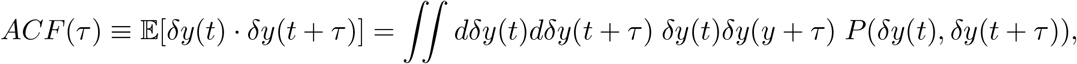

where 𝔼 [*X*] is the expectation value of *X*, δ*y*(*t*)= *y*(*t*) *-* 𝔼[*y*(*t*)], and

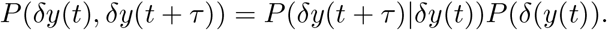

A static signal will stay perfectly correlated over time. However, as a dynamic signal *y*(*t*) evolves through time, (*e.g*., as it transitions between states), it becomes less correlated across longer time intervals. Consequently, the correlation captured by the ACF decreases as a function of τ. Because of this property, the ACF of an HMM reports simultaneously on the dynamic range of the signal, the noise in the signal, and the overall rates of transitions between states. Since HMMs are a well-defined probability model, the characteristic ACF for the process defined by an HMM can be calculated directly from the model parameters themselves [8]. For a discrete-time *K*-state HMM, the ACF is

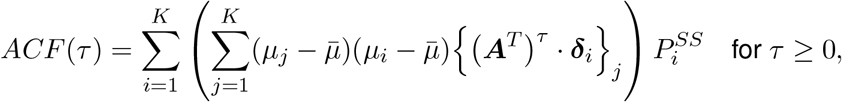

where μ_*i*_, σ_*i*_, ***A***, and 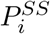 are defined as above, ***A***^T^ is the transpose of ***A***, δ_*i*_ is a column vector where the only non-zero element is a 1 in the i^th^ row, the notation *{****V*** *}*_j_ signifies the element in the j^th^ row of the column vector ***V***, and 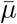 is the signal mean for the HMM at the steady state, given by 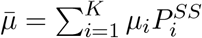.

To compare the results of an HMM modeling a simulated dataset with an inferred HMM that may or may not match the simulated HMM model, we calculate the ACF of the inferred HMM and then compare it to the ‘ground truth’ ACF calculated using the HMM parameters used in the simulation of the data. In our calculations, the limit lim _τ → ∞_ ***A***^*τ*^ for 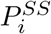 is approximated by exponentiating ***A*** to an arbitrarily large number — in our case, 100 times the inverse of the minimum *P*_*ij*_ in a specific transition matrix ***A***. Using this approach, we can generate the analytical ACF for any given HMM and circumvent complications in calculating the empirical ACF from specific instances of Markov chains inferred from time-limited trajectories (using, for example, the Viterbi algorithm [9]). The use of analytical ACFs ensures that any differences highlighted between HMM ACFs can be attributed to discrepancies in the model parameters themselves and thereby report on the abilities of specific inferred HMMs to explain the complete dynamics of a mesoscopic ensemble.

### 4 Kinetic Modeling with Composite HMMs

In the composite modeling approach, an HMM is inferred for each individual time-series before using a Gaussian mixture model (GMM) to cluster these results and provide ensemble-level information. The individual HMM for each trajectory is inferred using the variational Bayesian inference algorithm as implemented in the tMAVEN software package. The priors and likelihoods for the inference may be found in [10]. During this process, each trajectory is modeled with HMMs possessing a variable number of emission states (*K* = 1 to *K* = *K*_*ens*_, where *K*_*ens*_ is the number of states for the ensemble model, which in our case is either 2 or 4), and the HMM with the number of states that maximizes the evidence lower bound (ELBO) is chosen [10]. To identify the ensemble-level emission states, the composite approach then uses a *K*_*ens*_-state GMM to cluster the individual trajectory-level emission states, using prior distributions and likelihoods as specified in [11]. The GMM is inferred using a variational Bayesian algorithm as implemented in the tMAVEN software package. The values of the prior distribution hyper-parameters for the GMM means and precisions (which are the inverse of the squares of the standard deviations) are the same as the ones for the emissions of the HMM. Similarly, the priors used for the fractions of the GMM follow those used for the HMM initial probabilities [10]. The GMM takes as its input the set of individual idealized Viterbi paths [9] from each trajectory and is initialized with a K-means algorithm [11] with *K* = *K*_*ens*_.

In addition to the ensemble emission distributions, the GMM yields the ‘responsibilities’ of each state (*P*(*m* |*i*)), which are the probabilities that an individual trajectory-level emission state i corresponds to a specific ensemble emission state m. This allowed us to map the parameters for the individual trajectory-level HMM into the ensemble-level composite HMM. For example, the composite counts matrix ***𝒞***, which represents the number of transitions between all of the states in the dataset and thus may be thought of as the un-normalized form of the composite transition matrix ***𝒜***, can be assembled by adding the transition counts between the states for each trajectory, which is given by the individual counts matrix ***C***^*s*^ (for a given trajectory *s*). This, however, requires the mapping of the states between those that occur in the individual trajectory model to those that occur in the ensemble model. Using the corresponding responsibilities, thus, the transitions between the *m*^*th*^ and the *n*^*th*^ state of the ensemble may be estimated as

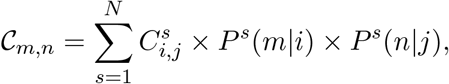

where the notation *B*_*ij*_ refers to the element on the *i*^*th*^ row and *j*^*th*^ column of the matrix ***B***, and *P*^*s*^ is the responsibility corresponding to the trajectory *s*. Having estimated the counts for all the states of the ensemble in this manner, ***𝒞*** is then row normalized to yield the transition matrix, ***𝒜***.

### 5 Kinetic Modeling with Global HMMs

The global modeling approach infers a single, ensemble-level HMM from a given set of experimental time series. This HMM is inferred using the variational Bayesian algorithm as implemented in the tMAVEN software package. The priors for the global HMM are the same as the individual HMM above and can be found in [10]. Since the trajectories are modeled as independent and identically distributed (i.i.d.) to one another, the global log-likelihood is the sum of the individual trajectory likelihoods given by

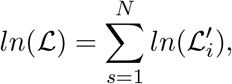

where 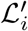 is the the HMM likelihood function for a single trajectory as reported in [10]. This is equivalent to concatenating the ensemble of trajectories into a single, long trajectory which is then analyzed with an HMM.

### 6 Estimation of Higher-level Transition Probabilities for hHMMs

In the above sections, we showed how we could relate the transition probabilities of the states at the production level (in our case, *d* = 3, Figure S3a) to the parameters of the hHMM. Because of the structure of the hHMM, these can be achieved for states at the higher levels (in our case, *d* = 2) of the HMM as well. These can be written as

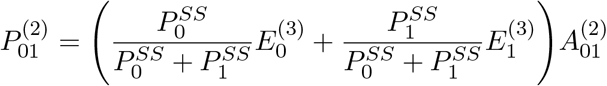

and

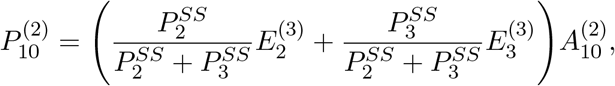

where the superscript (*n*) denotes the level the quantity refers to, the hHMM parameters are defined as above, and the steady-state probabilities for the production-level states 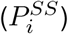 are calculated using the diagram method [6]. These expressions allow us to calculate the probabilities of transition between the slow (*s*) and fast (*f*) phases (*P*_*sf*_ and *P*_*fs*_) from the inferred hHMMs for our simulated datasets.

### 7 tMAVEN Software Organization

The tMAVEN software is written in Python and provided as open-source code. It consists of (*i*) a stand-alone graphics-less backend object that can run all of the analysis and that is available for scripting purposes; and (*ii*) the frontend graphical user interface (GUI) that facilitates interactive use of a backend instance. This section covers the use of the tMAVEN GUI and the design principles and organization of that software. Broadly tMAVEN is meant to enable users to easily (*i*) preprocess their data, (*ii*) perform kinetic modeling to analyze their data, (*iii*) reproducibly create publication-quality plots of their data and/or analysis. Additional documentation, including tutorials and example datasets, can be found on the tMAVEN documentation website (https://gonzalezbiophysicslab.github.io/tmaven/).

#### 7.1 Preprocessing data with tMAVEN

The main window of tMAVEN consists of a plot of a single trajectory (Fig. S4a); this is the _1_current_1_ trajectory. Users may use the keyboard arrow keys, a mouse scroll wheel, or the slider at the bottom of the GUI to rapidly scan through their trajectories. Additionally, the _1_Molecule Table_1_ dock provides information on each of the trajectories (Fig. S4a). Trajectories may be loaded into tMAVEN from a variety of sources (*e.g*., HDF5 files, ASCII, NumPy binary, *etc*.). Newly loaded trajectories are added to the currently loaded trajectories, rather than replacing them; this allows large, combined datasets from multiple sources (*e.g*., different movies) to be easily created.

Trajectories may be sorted into numbered _1_classes_1_ using the number keys or by selecting groups of trajectories in the _1_Molecule Table_1_ and using menu options (*i.e*., Tools > Classes > Set Class). The suggested workflow is for a user to rapidly survey all trajectories and perform an initial classification (*e.g*., ‘good’ trajectories into class 1 while ‘bad’ trajectories stays as the default class 0). Trajectories may be removed using menu options (*e.g*., Tools > Cull > Cull Class). Importantly, however, a basic design principle of tMAVEN is that the raw trajectories are immutable and are never directly changed, even when saved (see below). This is important for maintaining accurate records; notably, the filename of the origin of every trajectory is recorded and propagated as meta-data in subsequently saved files (see below) to enable the user to track the ‘source’ of their data. Many of the preprocessing options in tMAVEN do affect the trajectory values, however, they do so by acting on a copy of the ‘raw’ data called ‘corrected.’ For example, if a user applies a filter (*e.g*., a moving average) or a correction (*e.g*., bleed-through) to their data, the action affects the ‘corrected’ trajectories, which maybe be reset back to the ‘raw’ trajectories at any time (*i.e*., Tools > Corrections > Reset). All plotting and analyses operate using the corrected trajectories, however, only the raw trajectories are ever saved. Thus, all corrections must be reapplied if tMAVEN is closed or the data is cleared (*i.e*., File > Clear Data) and reloaded.

Aside from enabling facile sorting of trajectories, tMAVEN preprocessing capabilities are largely concerned with identifying single sub-regions of each trajectory for analysis. This enables users to isolate the region of their trajectories where, *e.g*., fluorophore photobleaching has not affected the intensity values. Sub-regions are determined by setting a ‘pre’ and a ‘post’ time point (left or right clicking, respectively) for each trajectory. Automatic photobleaching detection methods can be used to process multiple trajectories at the same time. The design choice of only allowing a single sub-region *per* trajectory is meant to forcibly ensure users avoid improperly affecting the underlying kinetics by inducing strange conditional dependencies, *e.g*., by excluding regions of fluorophore blinking which may preferentially occur from certain intensity states. Recognizing that certain types of binding experiments benefit from isolating multiple binding events *per* trajectory, tMAVEN does allow users to ‘split’ a trajectory by cloning it and setting the post-point of the first trajectory to be the pre-point of the second trajectory (using the ‘c’ key). In this manner, isolating multiple regions *per* trajectory can be performed by creating multiple copies of the trajectory and using the single sub-region of each trajectory to isolate separate regions of the trajectory. Note that such split trajectories are tracked by tMAVEN and can be recombined back into a single trajectory (using the ‘v’ key).

Trajectories in tMAVEN are preferentially saved in a variation of the Single-Molecule Dataset (SMD) format [12] but may also be exported in ASCII or NumPy binary format. This variation is primarily to enable use of HDF5 files, rather than the original JSON format, which enables more facile cross-program compatibility as HDF5 datasets self-describe data formats and libraries for parsing HDF5 files exist for many programming languages. tMAVEN considers each dataset saved in SMD format as an HDF5 group; thus, multiple SMD datasets may be saved in the same HDF5 file (Fig. S4b). An example HDF5 file with two real single-molecule fluorescence resonance energy transfer (smFRET) SMD datasets is available on the tMAVEN documentation website at https://gonzalezbiophysicslab.github.io/tmaven/examples/example_smd.hdf5. Notably, all information relating to tMAVEN preprocessing (*e.g*., class assignments, pre- and post-points) are not directly specified as a requirement of the SMD format, and so are instead saved as an HDF5 group called tMAVEN within the SMD group (Fig. S4b). The SMD format saves the ‘raw’ trajectories as a dense array of data in the order: number of trajectories, time, and then ‘color’ channels. Also included for each trajectory is a ‘source index’, which specifies the ‘source’ of that trajectory. To ensure proper tracking of trajectory data origins, meta-data about these sources are included in the ‘sources’ group within the SMD group (Fig. S4b).

#### 7.2 Kinetic Analysis with tMAVEN

Once a user has obtained a pre-processed set of trajectories, they may analyze these trajectories using a kinetic model. The design principles of tMAVEN are to consider (*i*) every initialized instance of a kinetic model as a ‘model’ (*e.g*., the untrained instance of an HMM is a different model than the trained version of that HMM), and (*ii*) models as representing the kinetics of macroscopic ensembles of molecules. Thus, while a user may ‘update’ a model by inferring kinetic parameters from a single or even several trajectories, the updated model technically describes an entire ensemble of hypothetical molecules. This means that all parameters of a model exist outside of the context of the particular trajectories used to estimate that model. In order for a user to _1_see the model of their data_1_, the model must be projected onto those trajectories, a process we call ‘idealization’. Note, this means that any trajectory can be idealized with a model, regardless of whether or not that trajectory was involved in estimating the model in the first place.

tMAVEN includes the ability to create several types of kinetic models that have been specialized for sm-FRET data (*e.g*., vbFRET [10]), however, these may be adapted and/or new types may be added. There are only a few requirements for a ‘model’ instance in tMAVEN. Currently these are limited to the parameters required to represent a Gaussian-distributed mixture model that can be overlaid onto a histogram of the trajectory values (*n.b*., this is a loose requirement required only for certain plots), and an ‘idealize’ function that takes an input trajectory and returns the idealized version of that trajectory (Fig. S4c). These idealized trajectories are often ‘Viterbi paths’, and are displayed overlaid on top of the original trajectory in the main tMAVEN window.

Notably, tMAVEN enables users to estimate multiple different models of their data. These are dynamically held in the *maven.modeler* object in a list called *models* (Fig. S4d). Only one model may be the ‘active’ model at a time, however, the user may change the active model (Modeling > Change Active), remove models (Modeling > Remove Models), or load and save models (Modeling > Load Active/Export Active). Dwell time analyses, based upon the state-space idealization of the loaded trajectories using a model, can be also performed (*i.e*., Modeling > Analyze Dwell Times). Additionally, several of the plotting options will include additional features, such as post-synchronization, based upon these state-space idealizations (*e.g*., FRET HIST 2D; see below). Finally, information about each model, such as kinetic parameters, may be obtained as a ‘report’ (Modeling > Generate Report) that may be viewed in the tMAVEN log (View > Show Log).

#### 7.3 Data Visualization with tMAVEN

While there are several types of plots that may be made in tMAVEN, they all have a common design principle of enabling the user to programmatically customize all visual aspects in order to obtain complete continuity between plots across multiple datasets. These customizations are performed by changing the plot preferences in the Preferences dock (*e.g*., *‘plot.intensity_max’* changes the y-axis maximum value); note that each plot has its own set of preferences. Once a user settles on a set of preference values for a plot, they may create a script to programmatically set all those values in one step rather than editing the underlying tMAVEN code.

A plot of the ‘current’ trajectory is shown in the main interface (Fig. S4a). This may be saved by clicking on the ‘disk’ icon in the toolbar underneath the plot; suggested file types are .pdf and .png. The analysis plots that are accessible under the ‘Plots’ menu bar include data from all trajectories. The ‘FRET Hist 1D’ plot is a one-dimensional histogram of the normalized data values (*i.e*., E_FRET_) between the pre- and post-points for all the trajectories; if there is an active model, a Gaussian mixture representing the states of that model are shown overlaid on this histogram. The ‘FRET Hist 2D’ plot is a two-dimensional heat map of the E_FRET_ values at each time point; if there is an active model, this histogram may be post-synchronized to a specified transition between two (or more) of the identified states. The ‘FRET TDP’ plot is a transition density plot of the E_FRET_ values between neighboring datapoints (*i.e*., initial and final E_FRET_); it defaults to a log-scaled colorbar. The ‘Dwell Times’ plot creates survival plots of a dwell time analysis (Modeling > Analyze Dwell Times) that can be fit and plotted along with the residual. Finally, custom plots are readily created by writing scripts (see the tMAVEN documentation website for examples of creating a ‘rastergram’ or plotting stochastic entropy production *vs*. time).

## Supplemental Figures

**Figure S1:**
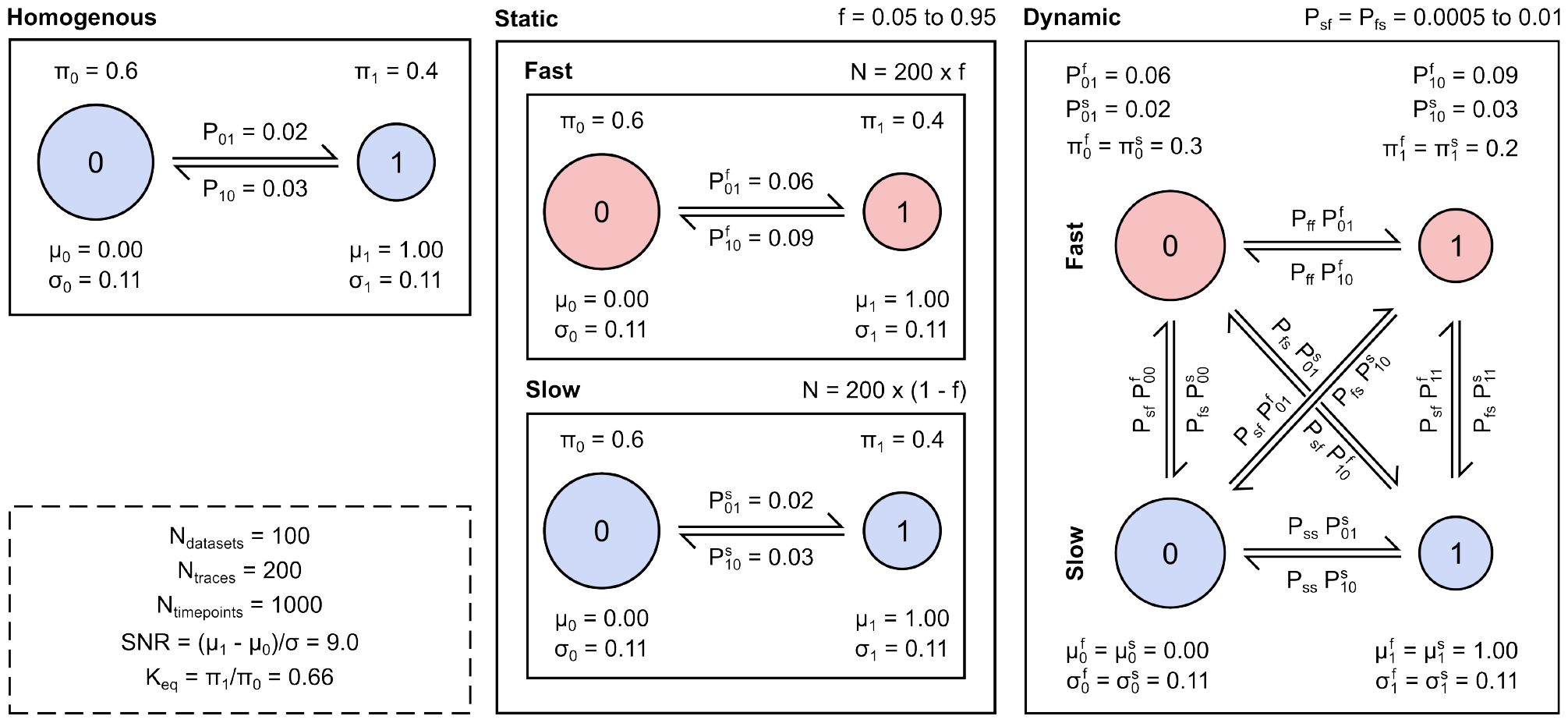
Parameter details for the simulated datasets. The underlying parameters for the three simulated datasets with Markovian kinetics (self-transition probabilities are omitted) and Gaussian emission distributions: *(left)* two-state Markovian ensembles, where the states are 0 and 1 (Homogeneous), *(center)* ensembles of varying proportions of two, non-interconverting two-state subpopulations exhibiting fast and slow kinetics (Static), and *(right)* ensembles of systems representing increasing transitions between a ‘slow’ phase (s) and a ‘fast’ phase (f) of the two emission states (Dynamic). The common parameters used for these datasets are shown *(bottom left)*.

**Figure S2:**
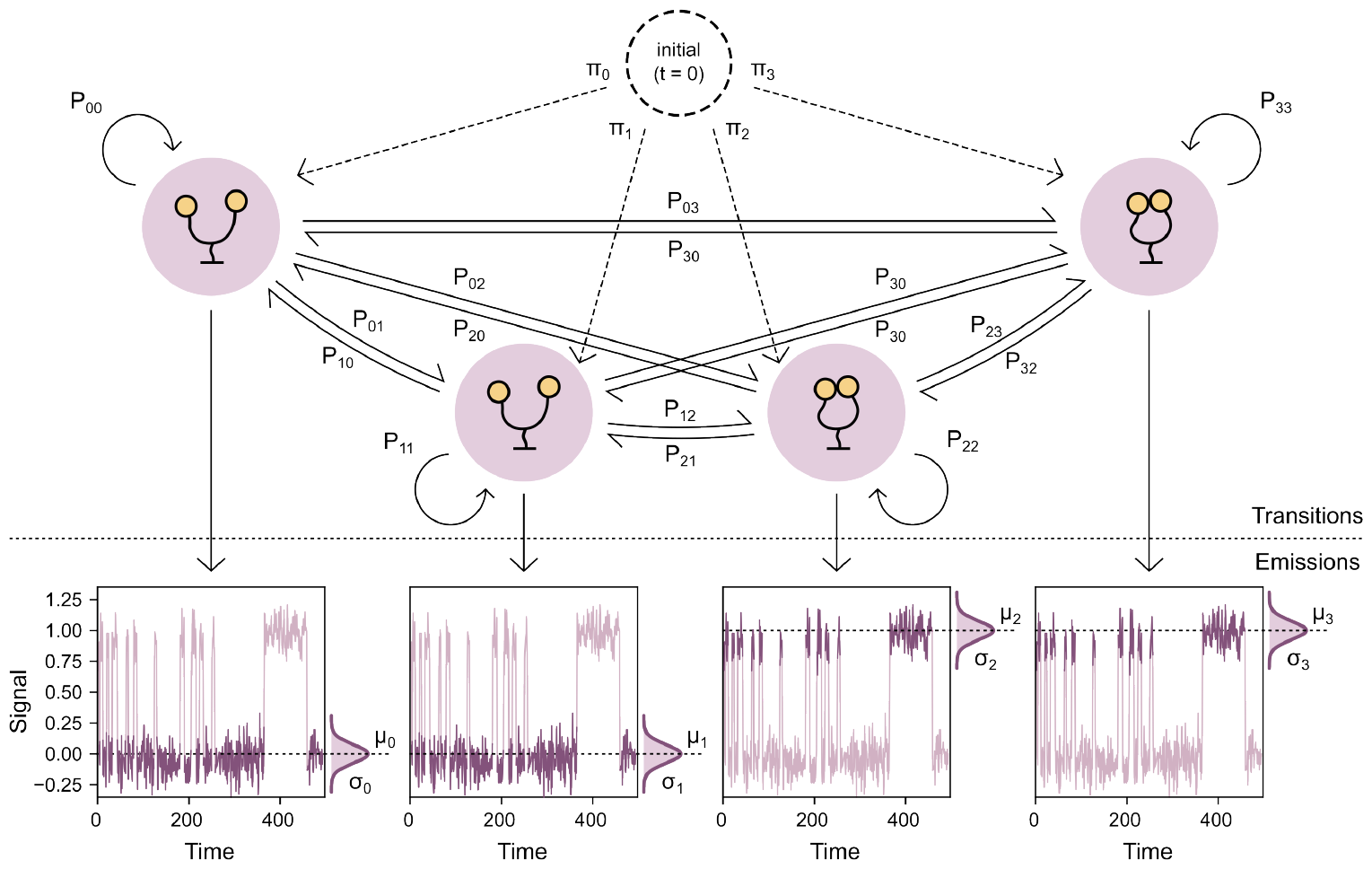
Schematic diagram of a four-state HMM. Diagram of the tree structure for a four-state HMM showing the transition and the emission parameters.

**Figure S3:**
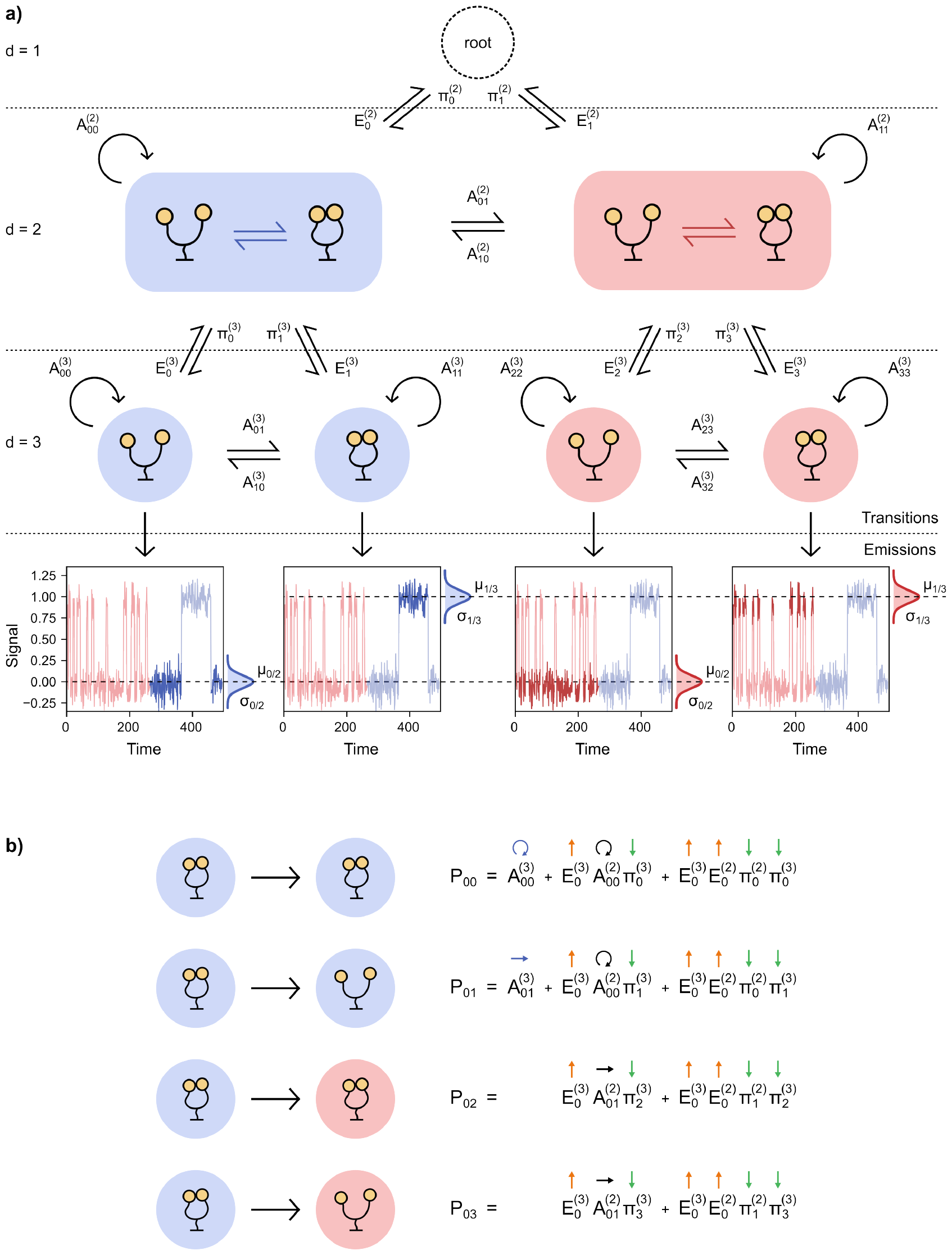
Schematic diagram of two-level four-state hHMM. **a)** Diagram of the tree structure for a two-level, four-state hHMM showing the transition and emission parameters. **b)** Transition probabilities from the production-level state 0 to the other production-level states decomposed into contributions from different pathways that traverse the tree structure to reach the final state.

**Figure S4:**
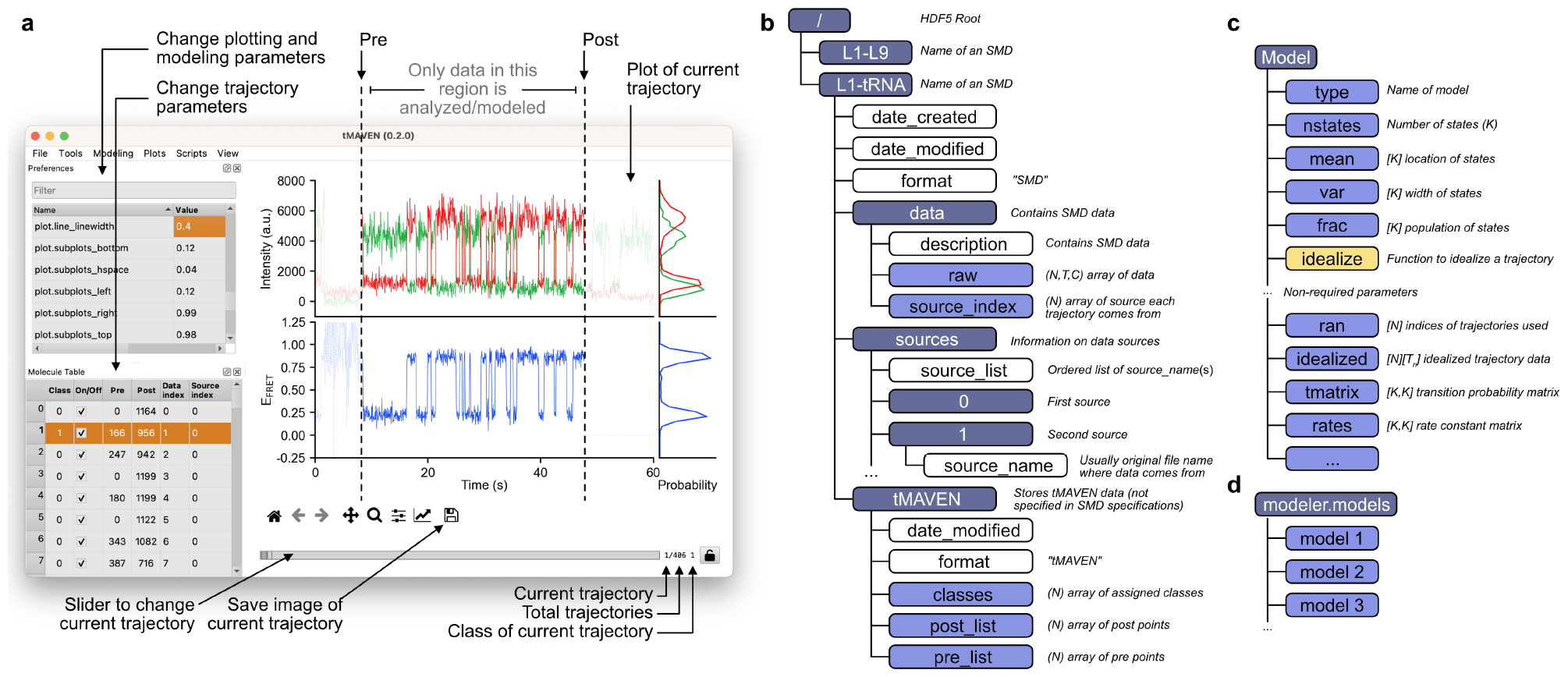
Annotated tMAVEN interface and data structures. a) The user interface for tMAVEN is shown. The Preferences dock in the upper-left allows users to change any parameters involved in plotting, analysis, *etc*.. The Molecule Table dock in the lower-left provides information about status of the trajectories and can be used to select multiple trajectories at once. b) Schematic of two example datasets saved as individual Single-Molecule Datasets (SMD) in a single HDF5 file. HDF5 group names are shown in dark blue, HDF5 attributes are shown in white, and HDF5 datasets are shown in light blue. c) A schematic of a kinetic model Python object in tMAVEN with the parameters required for all types of kinetic models on top, and several non-required model-specific parameters shown on bottom. d) After being estimated, every kinetic model is stored in tMAVEN’s modeler object under the models list. The currently active model may be chosen from amongst these models.

**Figure S5:**
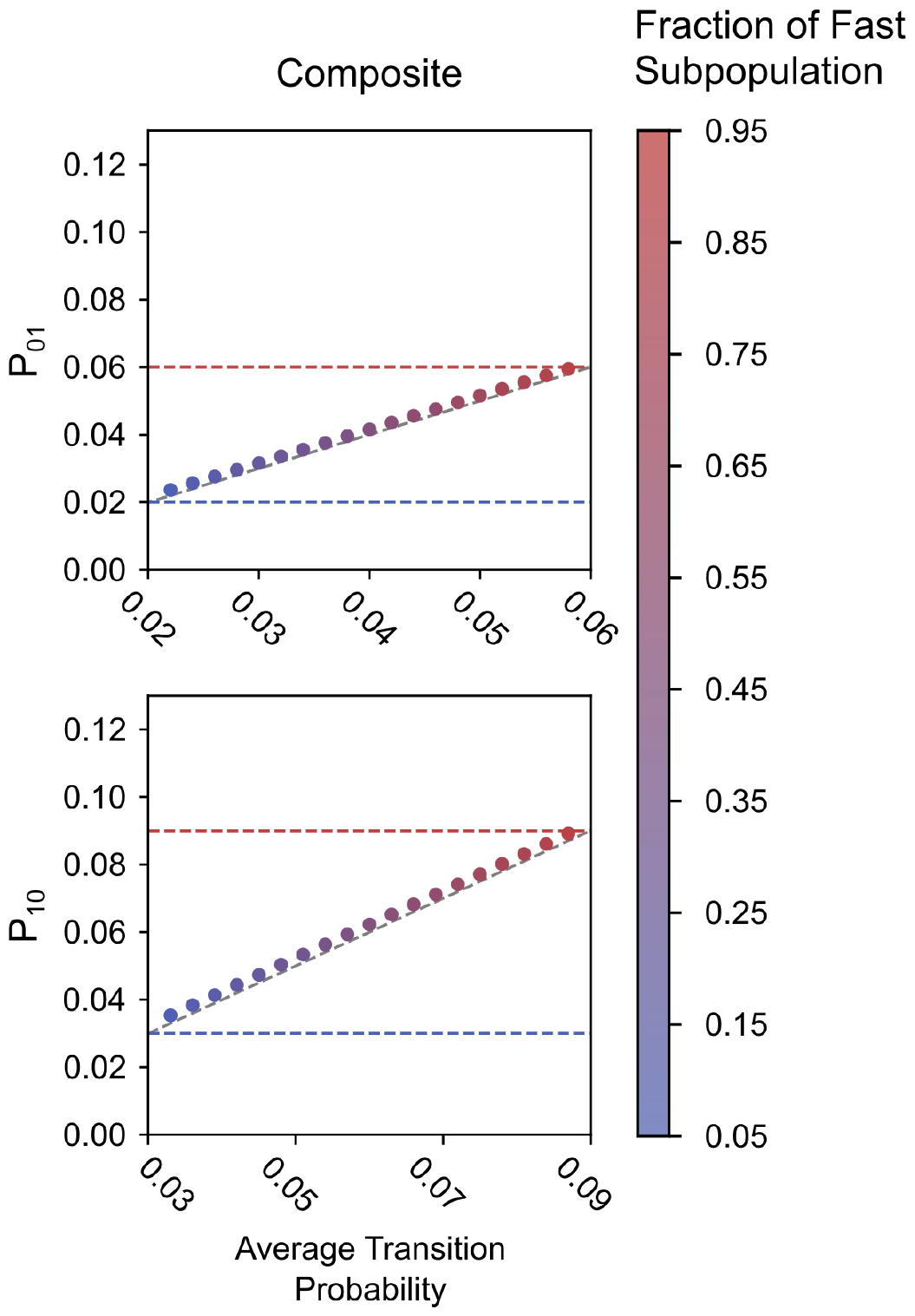
The effect of static heterogeneity on kinetic modeling using composite HMMs. The ensemble-level transition probabilities for the observed states inferred using composite HMMs as a function of the average transition probability of the observed states (calculated using the proportions of fast- and slow-transitioning molecules). The dashed grey line denotes identity. The dashed red and blue lines denote the transition probabilities from each state for the subpopulation of fast- and slow-transitioning molecules respectively.

**Figure S6:**
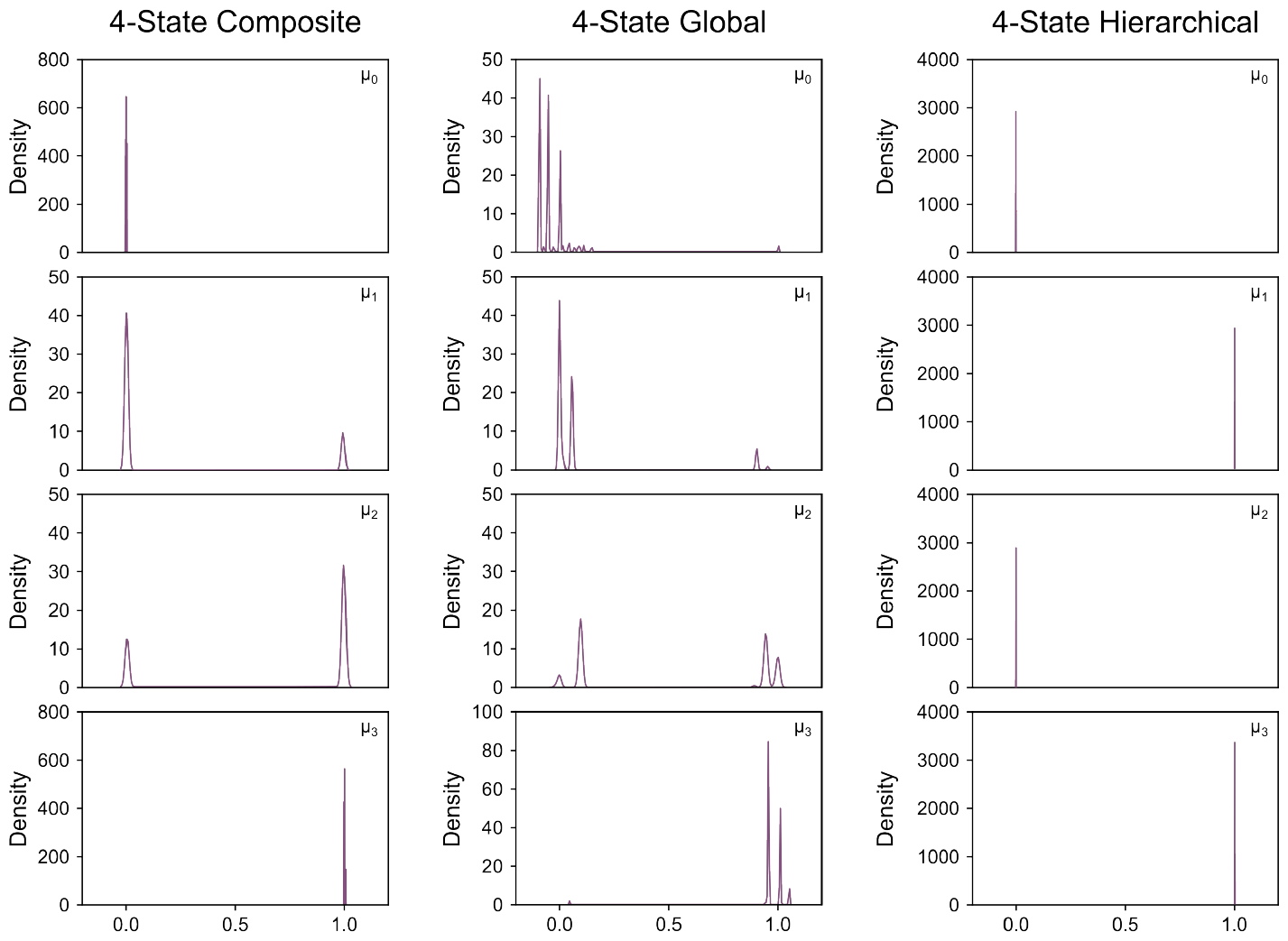
Effect of static heterogeneity on emission distributions for different four-state kinetic models. Kernel density estimated distributions of the emission means for the four states inferred using composite HMMs, global HMMs, and two-level hHMMs for a statically heterogeneous ensemble where the fraction of the fast subpopulation is 50%.

**Figure S7:**
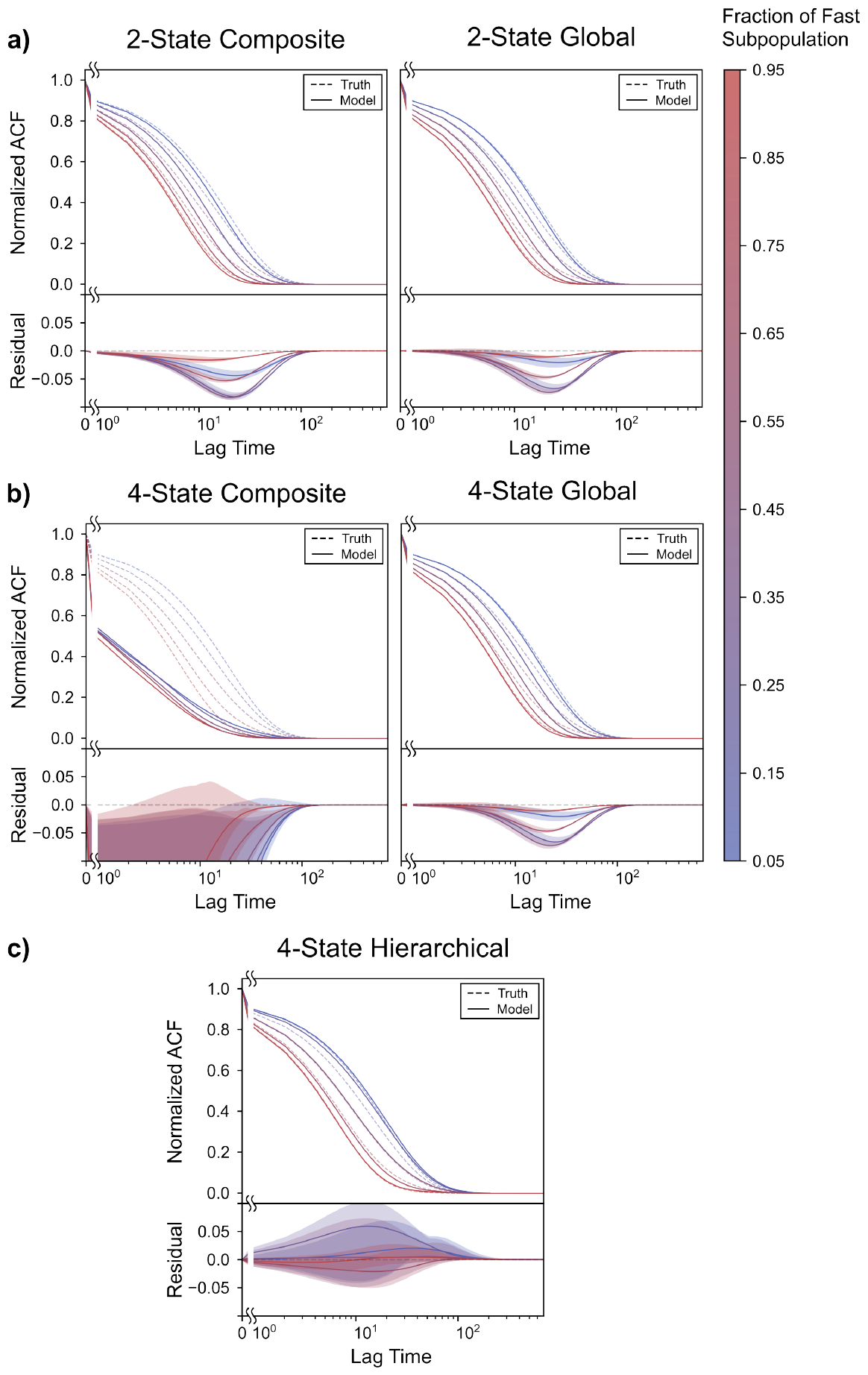
Comparison of ACFs for varying levels of static heterogeneity analyzed using different kinetic models. **(a)** *(top)* The true ACF (dashed) along with the mean of the inferred ACFs (solid) for Static datasets where the fraction of the fast subpopulation was 5%, 25%, 50%, 75%, and 95%, analyzed using two-state composite *(left)* and global *(right)* HMMs, along with *(bottom)* the corresponding mean (solid) of the residuals of the inferred ACFs to the true ACF. **(b)** *(top)* The true ACF (dashed) along with the mean of the inferred ACFs (solid) for Static datasets where the fraction of the fast subpopulation was 5%, 25%, 50%, 75%, and 95%, analyzed using four-state composite *(left)* and global *(right)* HMMs, along with *(bottom)* the corresponding mean (solid) of the residuals of the inferred ACFs to the true ACF. **(c)** *(top)* The true ACF (dashed) along with the mean of the inferred ACFs (solid) for Static datasets where the fraction of the fast subpopulation was 5%, 25%, 50%, 75%, and 95%, analyzed using two-level, four-state hHMMs, along with *(bottom)* the corresponding mean (solid) of the residuals of the inferred ACFs to the true ACF. The shaded area denotes the region one standard deviation away from the mean. The dashed grey line corresponds to zero.

**Figure S8:**
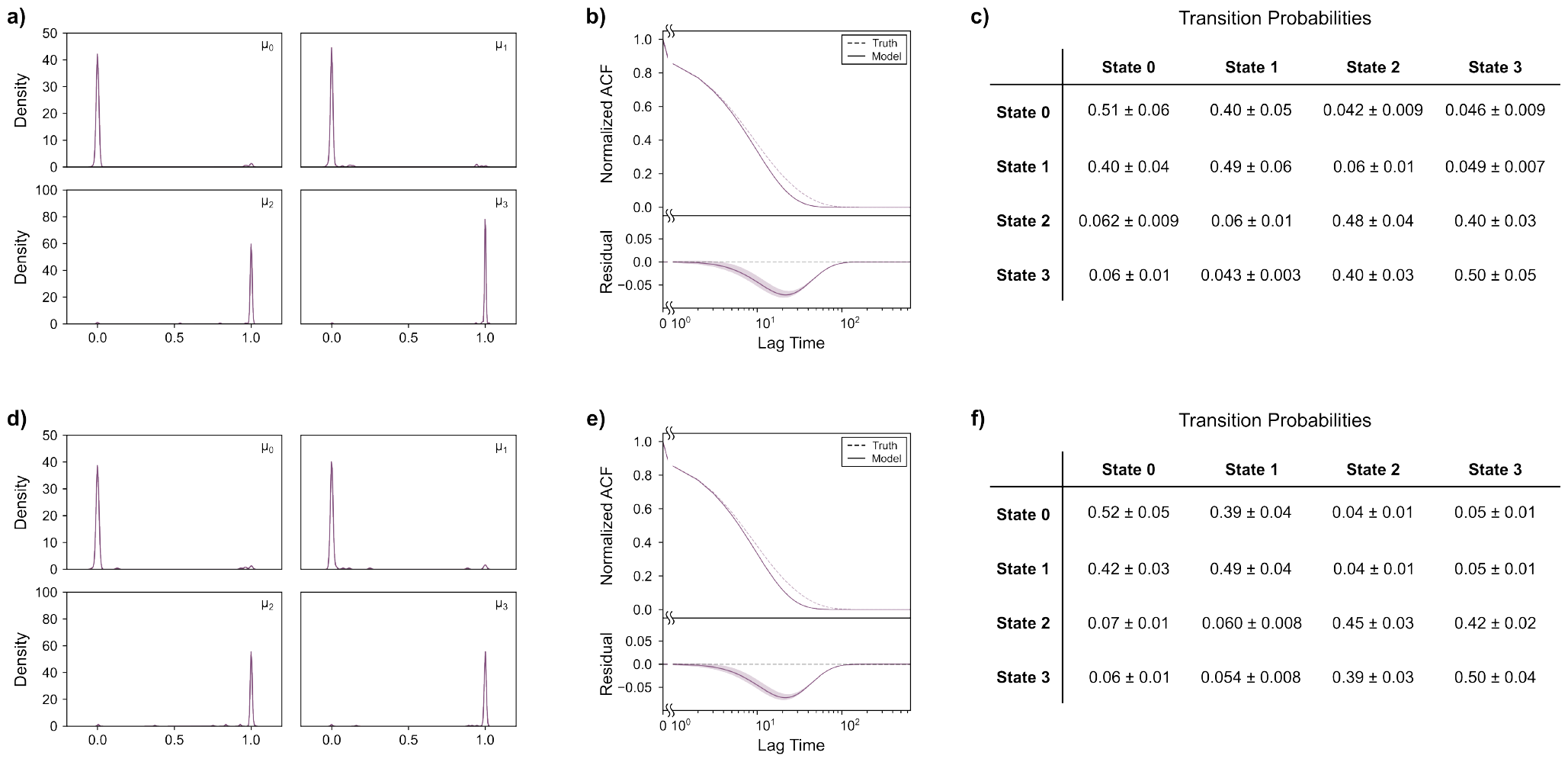
Effect of initializations at true parameter values on the modeling of static heterogeneity. **(a)** Kernel density estimated distributions of the emission means for the four states inferred using global HMMs for a statically heterogeneous ensemble where the fraction of the fast subpopulation is 50%, when the inference process is initialized at the true emission means, emission noise, and steady-state probabilities. **(b)** *(top)* The corresponding true ACF (dashed) along with the mean of the inferred ACFs (solid), along with *(bottom)* the corresponding mean (solid) of the residuals of the inferred ACFs to the true ACF. **(c)** The corresponding mean transition probabilities between the states inferred by the global HMMs. The errors are standard deviations of the probabilities across the 100 simulated ensembles. **(d)** Kernel density estimated distributions of the emission means for the four states inferred using global HMMs for a statically heterogeneous ensemble where the fraction of the fast subpopulation is 50%, when the inference process is initialized at the true emission means, emission noise, steady-state probabilities, and transition matrix. **(e)** *(top)* The corresponding true ACF (dashed) along with the mean of the inferred ACFs (solid), along with *(bottom)* the corresponding mean (solid) of the residuals of the inferred ACFs to the true ACF. **(f)** The corresponding mean transition probabilities between the states inferred by the global HMMs. The errors are standard deviations of the probabilities across the 100 simulated ensembles.

**Figure S9:**
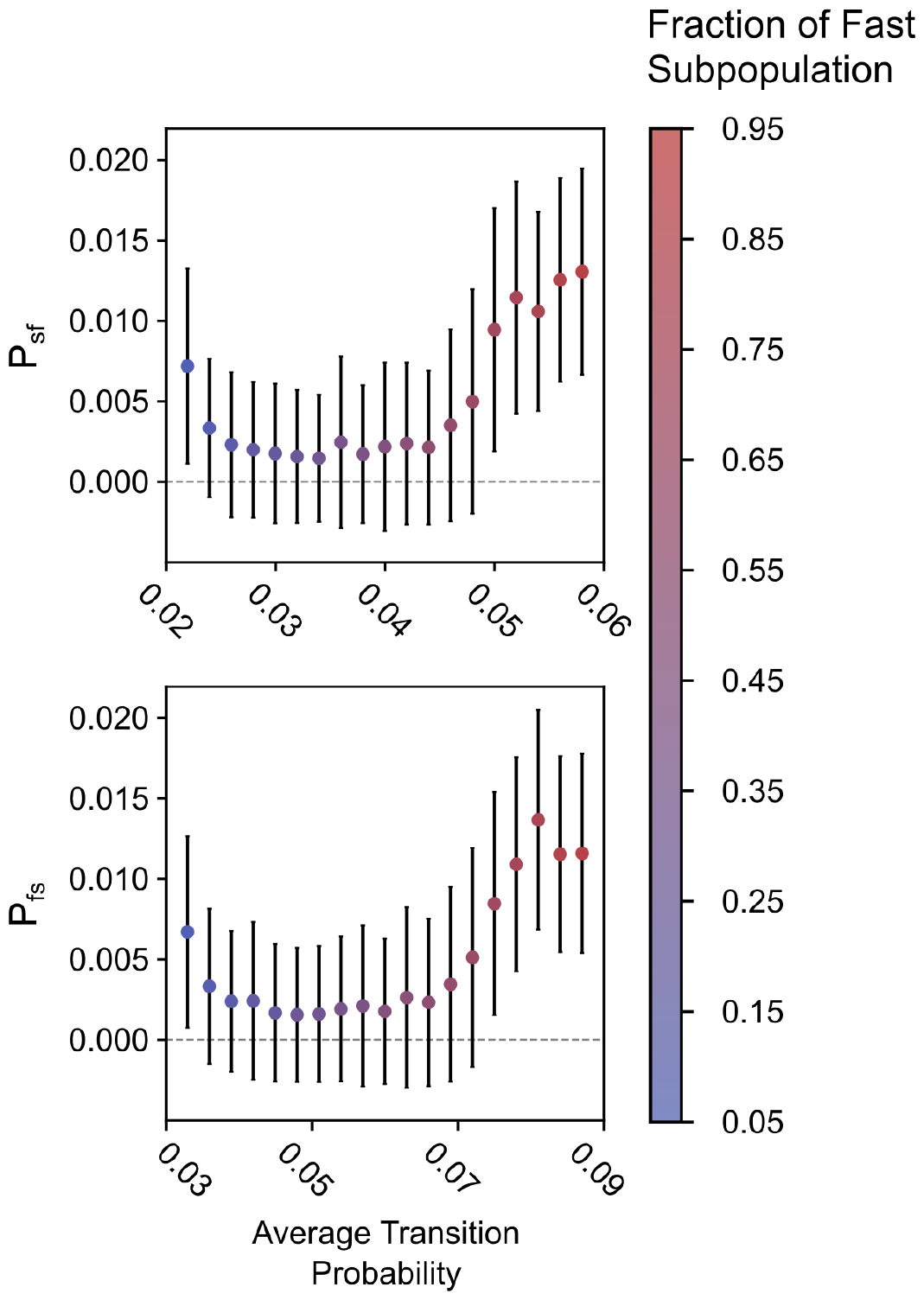
Hierarchical kinetics inferred using hHMMs for statically heterogeneous ensembles. The transition probabilities between the slow (s) and fast (f) subpopulations of Static datasets with varying levels of static heterogeneity, inferred using two-level, four-state hHMMs. The dashed grey line corresponds to the true value of transition probabilities between the two populations, which, in the case of static heterogeneity, is zero.

**Figure S10:**
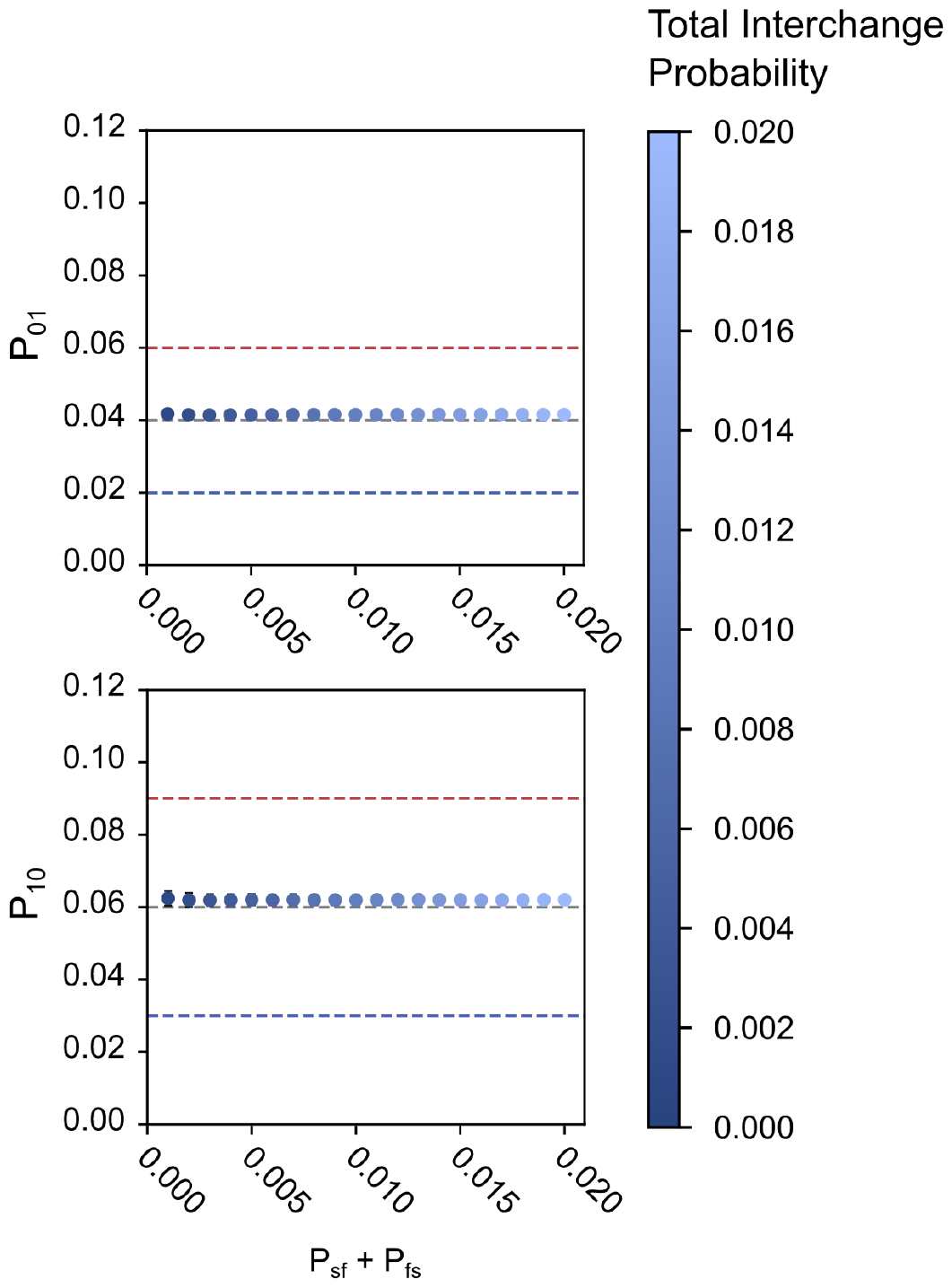
The effect of dynamic heterogeneity on kinetic modeling using composite HMMs. The ensemble-level transition probabilities for the observed states inferred using composite HMMs as a function of the total probability of transition between slow- and fast-transitioning phases (*P*_*sf*_ + *P*_*fs*_). The dashed red and blue lines denote the transition probabilities of each state for the fast- and slow-transitioning phases respectively. The dashed grey line denotes the ensemble average transition probability of each observed state.

**Figure S11:**
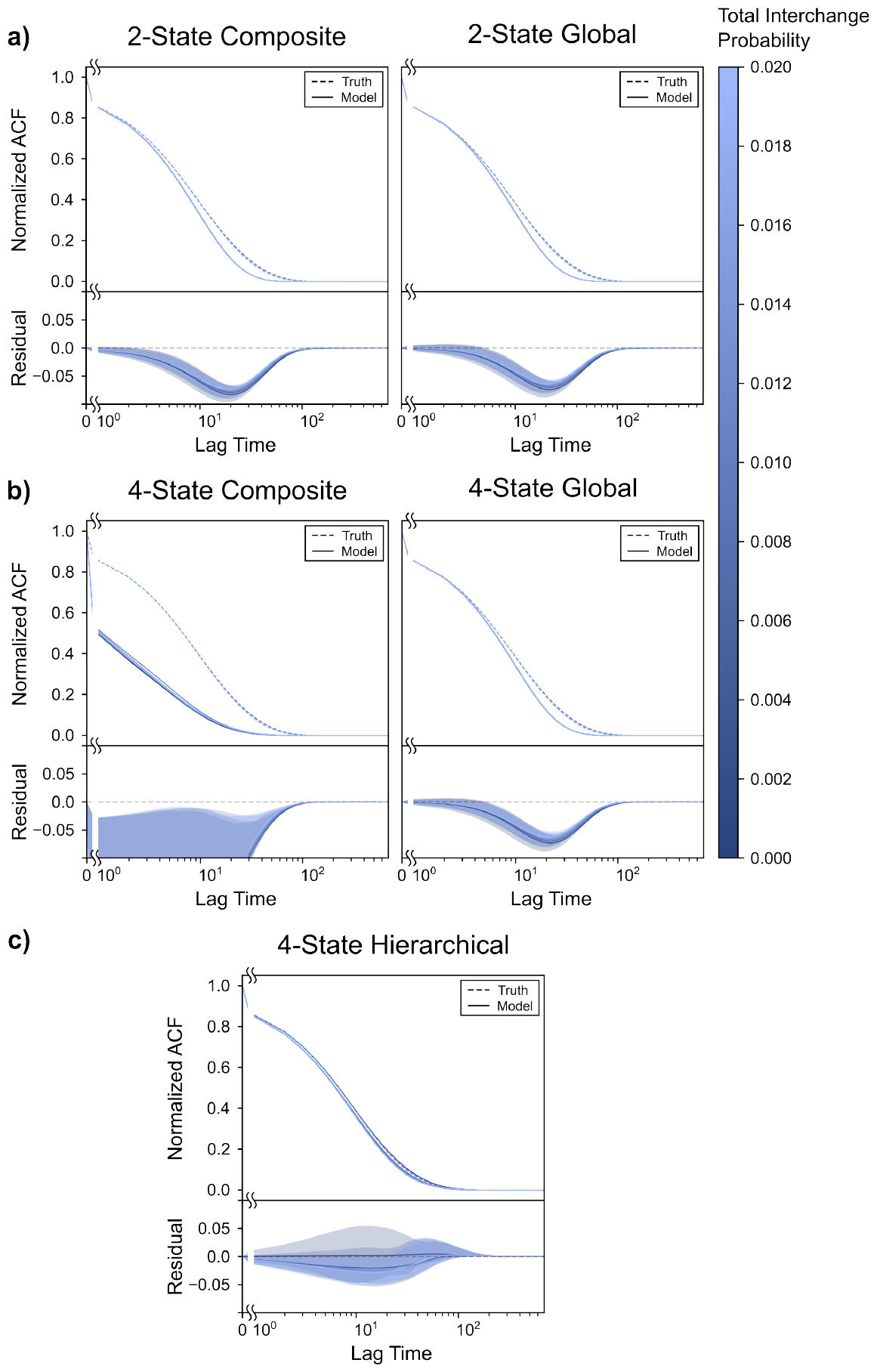
Comparison of ACFs for varying levels of dynamic heterogeneity analysed using different kinetic models. **(a)** *(top)* The true ACF (dashed) along with the mean of the inferred ACFs (solid) for Dynamic datasets where the total probability of interchange was 0.001, 0.005, 0.01, 0.015, and 0.02, analyzed using two-state composite *(left)* and global *(right)* HMMs, along with *(bottom)* the corresponding mean (solid) of the residuals of the inferred ACFs to the true ACF. **(b)** *(top)* The true ACF (dashed) along with the mean of the inferred ACFs (solid) for Dynamic datasets where the total probability of interchange was 0.001, 0.005, 0.01, 0.015, and 0.02, analyzed using four-state composite *(left)* and global *(right)* HMMs, along with *(bottom)* the corresponding mean (solid) of the residuals of the inferred ACFs to the true ACF. **(c)** *(top)* The true ACF (dashed) along with the mean of the inferred ACFs (solid) for Dynamic datasets where the total probability of interchange was 0.001, 0.005, 0.01, 0.015, and 0.02, analyzed using two-level, four-state hHMMs, along with *(bottom)* the corresponding mean (solid) of the residuals of the inferred ACFs to the true ACF. The shaded area denotes the region one standard deviation away from the mean. The dashed grey line corresponds to zero.

**Figure S12:**
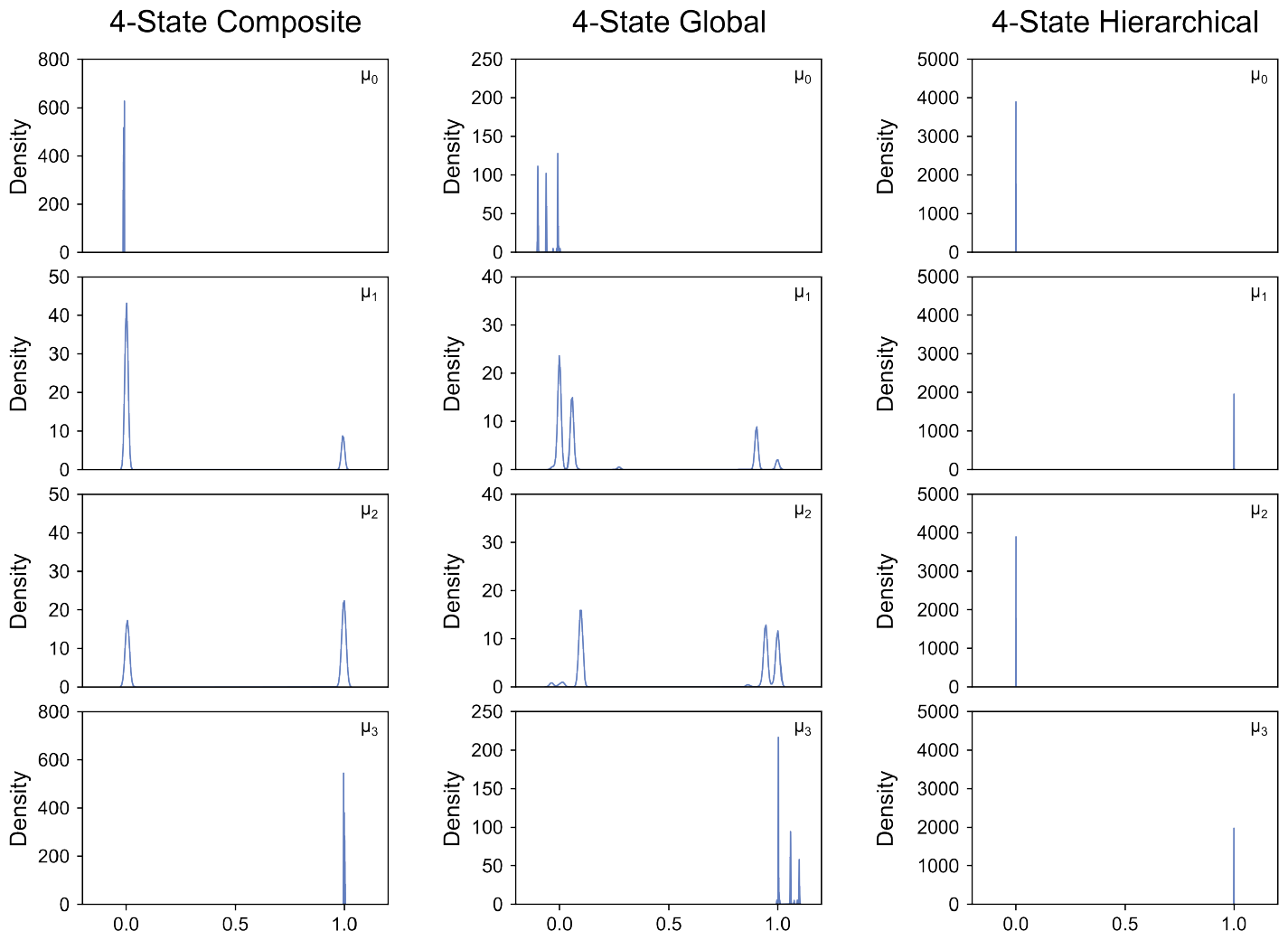
Effect of dynamic heterogeneity on emission distributions for different four-state kinetic models. Kernel density estimated distributions of the emission means for the four states inferred using composite HMMs, global HMMs, and two-level hHMMs for a dynamically heterogeneous ensemble where the total probability of interchange between the slow and the fast phases is 0.01.

**Figure S13:**
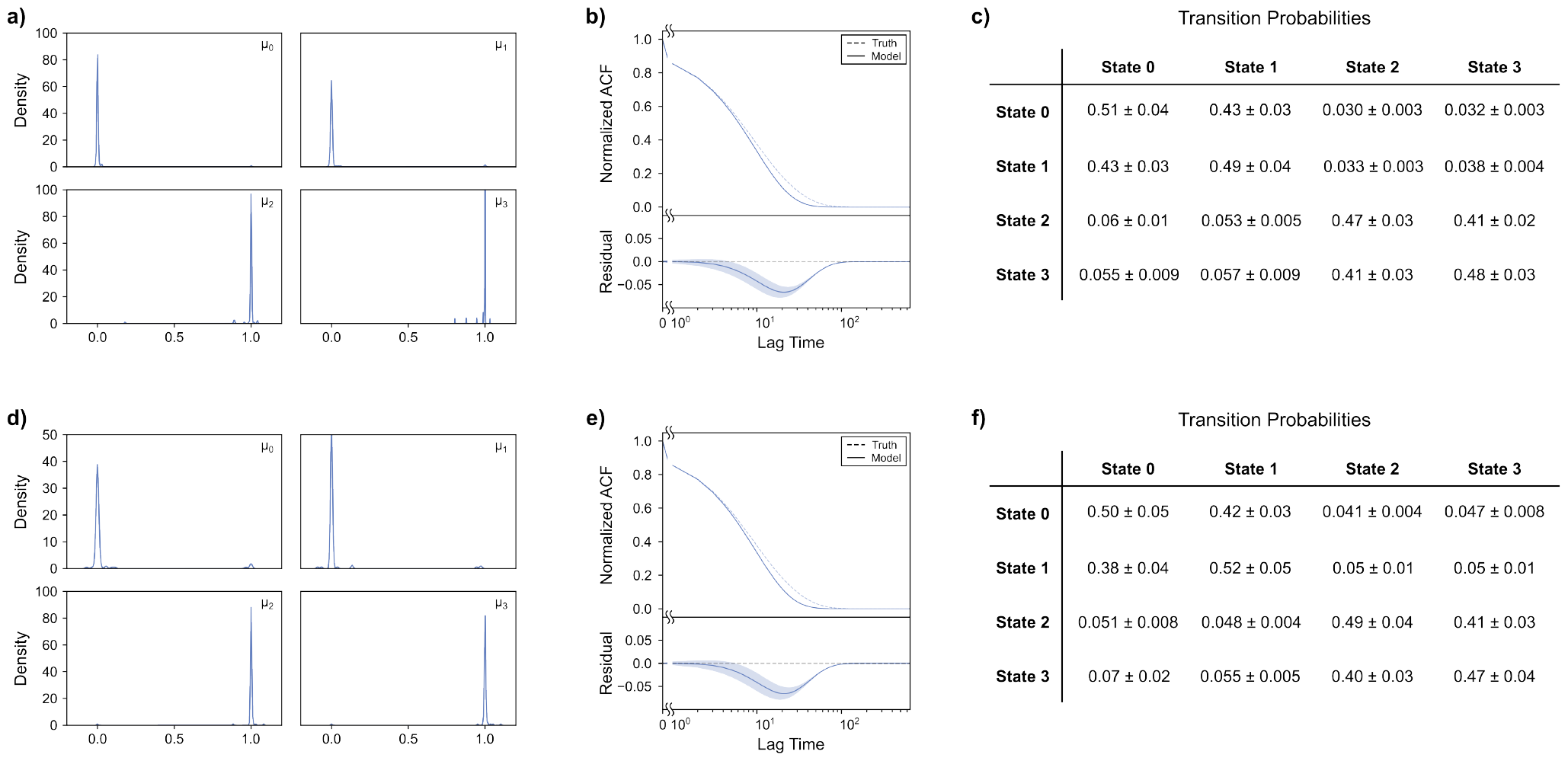
Effect of initializations at true parameter values on the modeling of dynamic heterogeneity. **(a)** Kernel density estimated distributions of the emission means for the four states inferred using global HMMs for a dynamically heterogeneous ensemble where the total probability of interchange between the slow and the fast phases is 0.01, when the inference process is initialized at the true emission means, emission noise, and steady-state probabilities. **(b)** *(top)* The corresponding true ACF (dashed) along with the mean of the inferred ACFs (solid), along with *(bottom)* the corresponding mean (solid) of the residuals of the inferred ACFs to the true ACF. **(c)** The corresponding mean transition probabilities between the states inferred by the global HMMs. The errors are standard deviations of the probabilities across the 100 simulated ensembles. **(d)** Kernel density estimated distributions of the emission means for the four states inferred using global HMMs for a dynamically heterogeneous ensemble where the total probability of interchange between the slow and the fast phases is 0.01, when the inference process is initialized at the true emission means, emission noise, steady-state probabilities, and transition matrix. **(e)** *(top)* The corresponding true ACF (dashed) along with the mean of the inferred ACFs (solid), along with *(bottom)* the corresponding mean (solid) of the residuals of the inferred ACFs to the true ACF. **(f)** The corresponding mean transition probabilities between the states inferred by the global HMMs. The errors are standard deviations of the probabilities across the 100 simulated ensembles.

**Figure S14:**
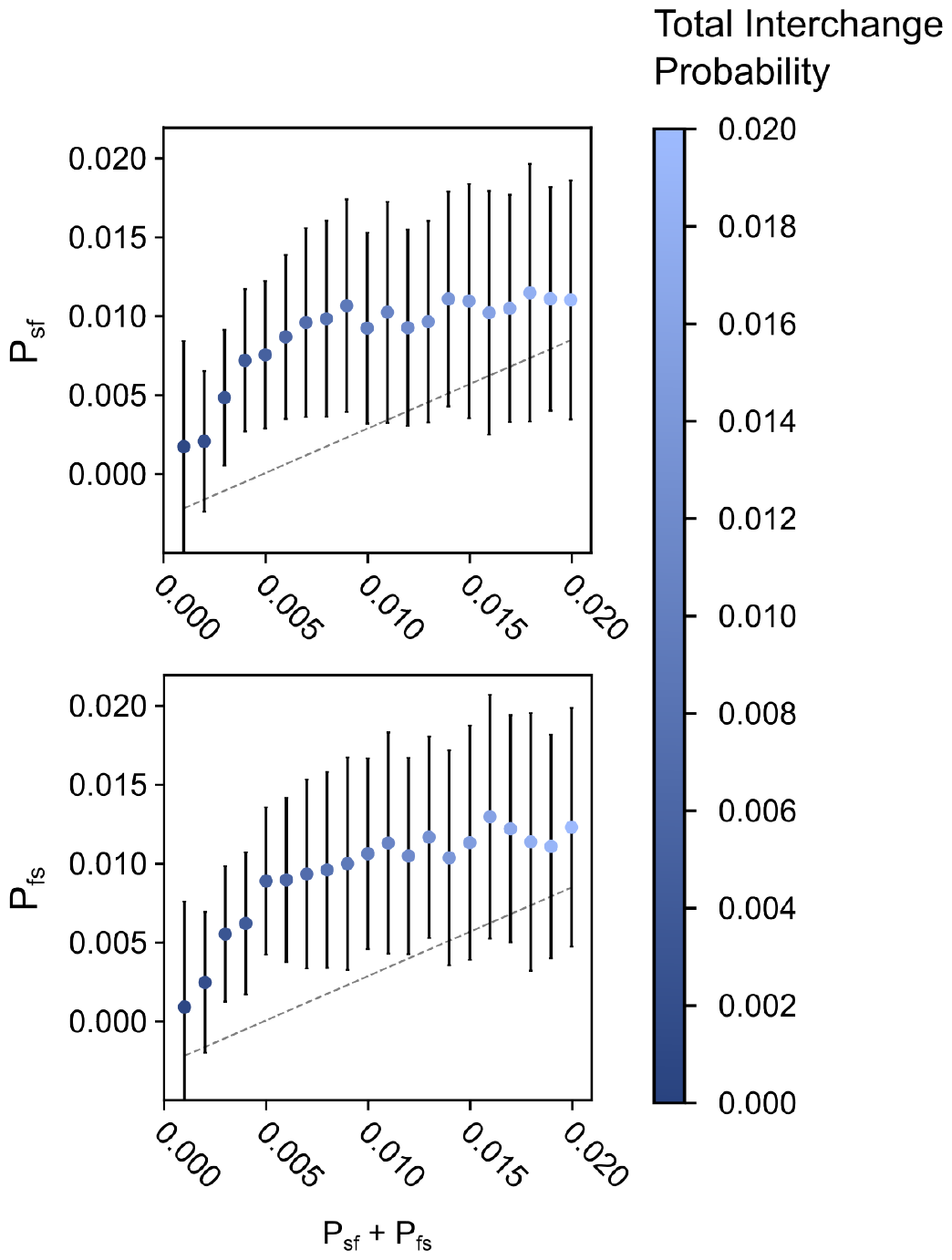
Hierarchical kinetics inferred using hHMMs for dynamically heterogeneous ensembles. The transition probabilities between the slow (s) and fast (f) subpopulations of Dynamic datasets with varying levels of dynamic heterogeneity, inferred using two-level four-state hHMMs. The dashed grey line corresponds to the true value of transition probabilities between the two populations.

